# Tandem Directional CTCF Sites Balance Protocadherin Promoter Usage

**DOI:** 10.1101/525543

**Authors:** Qiang Wu, Ya Guo, Yujia Lu, Jingwei Li, Yonghu Wu, Zhilian Jia

## Abstract

CTCF is a key insulator-binding protein and mammalian genomes contain numerous CTCF-binding sites (CBSs), many of which are organized in tandem arrays. Here we provide direct evidence that CBSs, if located between enhancers and promoters in the *Pcdhα* and *β*-globin clusters, function as an enhancer-blocking insulator by forming distinct directional chromatin loops, regardless whether enhancers contain CBS or not. Moreover, computational simulation and experimental capture revealed balanced promoter usage in cell populations and stochastic monoallelic expression in single cells by large arrays of tandem variable CBSs. Finally, gene expression levels are negatively correlated with CBS insulators located between enhancers and promoters on a genome-wide scale. Thus, single CBS insulators ensure proper enhancer insulation and promoter activation while tandem-arrayed CBS insulators determine balanced promoter usage. This finding has interesting implications on the role of topological insulators in 3D genome folding and developmental gene regulation.

## INTRODUCTION

Genetic studies have long described the phenomenon of Position Effect Variegation (PEV) [1], suggesting that the spatial organization of chromatin domains has an important influence on gene expression [2–4]. Early studies reveal that boundary elements, also known as insulators, restrict promoter activity from the position effects of its chromatin contexts [5, 6]. In particular, through a series of transgenic experiments, Grosveld and colleagues have identified dominant human *β*-globin flanking boundary elements which determine its position-independent expression in transgenic mice [5]. It has since been established that insulators play an essential role in shielding the position effects of chromatin conformation and in blocking enhancers or silencers from improperly activating or repressing non-cognate promoters, respectively [2, 3, 6–8].

The mammalian CTCF is the best characterized genome architectural protein that binds to insulator elements [2, 8]. CTCF directionally and dynamically binds to tens of thousands of CBSs (CTCF-binding sites) in mammalian genomes through the combinatorial usage of its 11 zinc fingers [9, 10]. CTCF, together with the associated Cohesin, a ring-shaped complex embracing DNA, mediates genome-wide long-range chromatin interactions [2]. Interestingly, these interactions are preferentially formed between forward-reverse CBS pairs [11–14]. The CBS elements in the boundaries between neighboring chromatin domains are configured in a reverse-forward orientation, which are thought to restrict enhancer activity to promoters within each chromatin domain [12, 15, 16]. Thus, the boundary CBS elements may function as insulators to block Cohesin loop extrusion [11, 12, 17–20]. However, whether and how internal CBS elements function as insulators remain incompletely understood.

Numerous studies have shown that CTCF/Cohesin-mediated chromatin loop domains or TADs (topologically associated domains) are important for gene regulation [4, 12, 15, 16, 21, 22]. Insertion, mutation, deletion, inversion, or duplication of CBS elements alters chromatin topology and gene expression [12, 14–16, 18, 22–24]. Emerging evidence suggests that spatial control of genome topology by CTCF/Cohesin regulates gene expression; however, how numerous CBS elements in mammalian genomes function as insulators to control proper promoter activation and balanced usage remains obscure.

## RESULTS

### Exogenous CTCF Sites as Protocadherin Insulators

Similar to the enormous diversity of DSCAM1 proteins in *Drosophila*, combinatorial *cis*- and *trans*-interactions between mammalian clustered cell-surface protocadherin (Pcdh) proteins, encoded by the three closely-linked gene clusters (*α, β*, and *γ*), endow individual neurons with a unique identity code and specific self-recognition module, which are required for neuronal migration, dendrite self-avoidance, and axon tiling in the brain [25–31]. The human *Pcdhα* cluster contains 13 highly-similar, tandem-arrayed, unusually-large “alternate” variable exons (*α1-α13*) and 2 divergent “ubiquitous” variable exons (αc1-αc2), followed by 3 downstream small constant exons (Figure 1A), reminiscent of the variable and constant genome organizations of immunoglobulin (Ig) and T-cell receptor (*Tcr*) clusters [25, 32]. Each of the 13 “alternate” variable exons (*α1-α13*) carries its own promoter, which is flanked by two forward-oriented CBS (*CSE* and *eCBS*) elements (Figure 1A). However, the *αc1* “ubiquitous” promoter carries only one forward-oriented CBS and the *αc2* promoter has no CBS element (Figure 1A). Two distal *Pcdhα* enhancers, *HS7* and *HS5-1*, are located downstream, and one of which, *HS5-1*, is flanked by two reverse-oriented CBS (*HS5-1a* and *HS5-1b*) elements [33, 34]. Multiple long-distance chromatin interactions between these enhancers and *Pcdhα* target promoters form a transcription hub and determine the promoter choice [34, 35]. We performed single-cell RNA-seq of mouse cortical neurons and found members of the *Pcdhα* cluster are expressed in single neurons in a combinatorial and stochastic manner (Figure 1B), similar to the stochastic monoallelic expression patterns of *Pcdhαin* single Purkinje cells in the cerebellum [27, 36].

**Figure 1.**
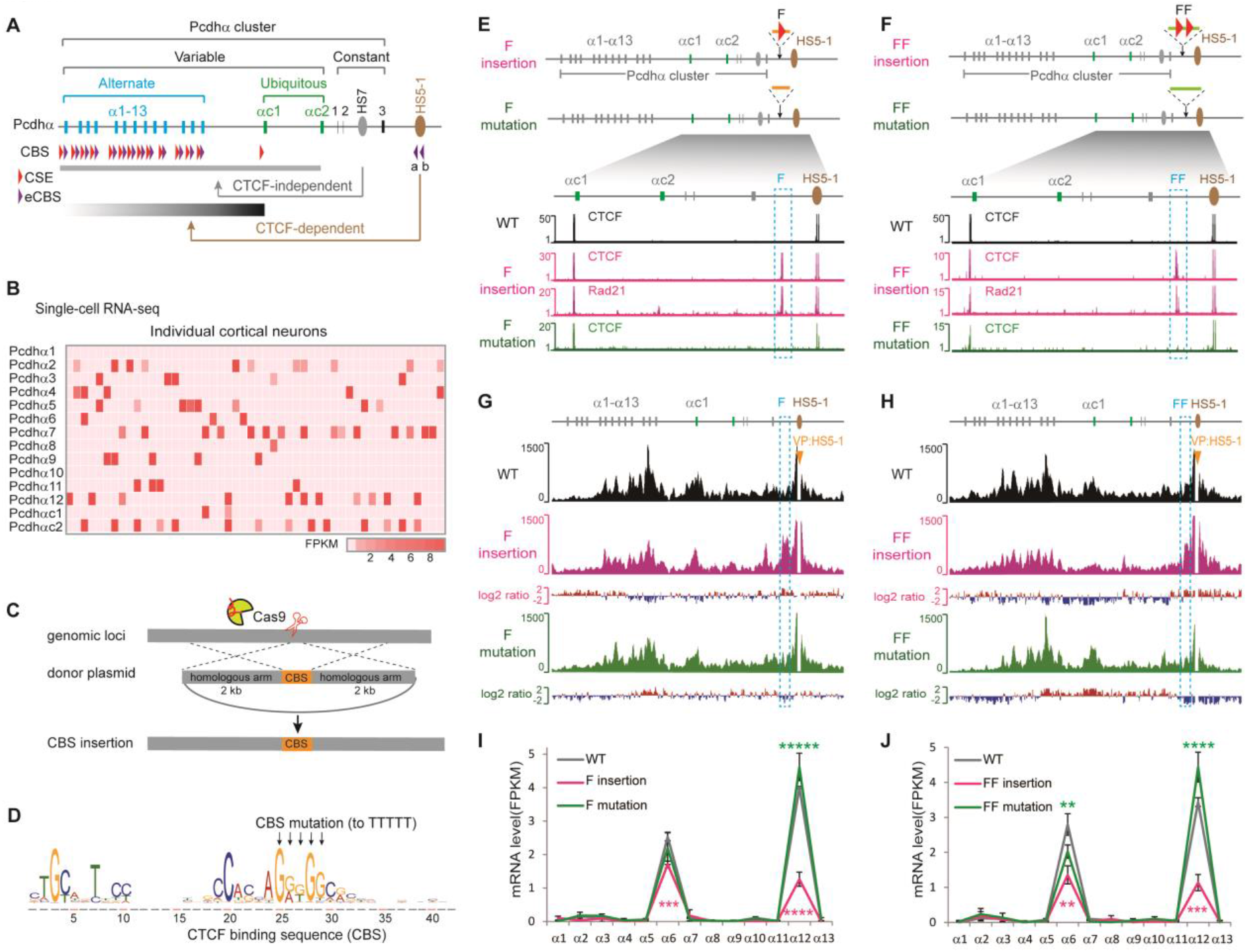
CTCF sites as insulators in the *Pcdhα* cluster through directional chromatin looping. (**A**) Schematic of the variable and constant *Pcdhα* genomic organization of the *Pcdhα* cluster. The variable region contains 13 alternate exons (*α1-α13*), each of which is preceded by its own promoter flanked by two forward CBS elements (indicated by tandem arrowheads), and two ubiquitous exons of *αc1* and *αc2*, of which only *αc1* contains a single CBS element. The constant region contains three exons which are spliced to each variable exon. Two distal enhancers, *HS7* (with no CBS) and *HS5-1* (flanked by two reverse CBSs), are located downstream. CBS: CTCF binding site; CSE: conserved sequence element; eCBS: exonic CBS. (**B**) Single-cell expression patterns of the *Pcdhα* genes in mouse neocortical cells. (**C**) CRISPR insertion of CBS elements by homologous recombination. (**D**) Schematic of CTCF binding motif and its mutation. (**E** and **F**) CTCF and Rad21 ChIP-seq of single-cell CRISPR clones with one or two forward-oriented CBSs inserted into the location between *Pcdhα* cluster and its downstream *HS5-1* enhancer. (**G** and **H**) QHR-4C with *HS5-1* as a viewpoint (VP, arrowheads). Log2 ratios (insertion vs wildtype or mutation vs insertion) are shown under the 4C profiles. (**I** and **J**) Gene expression levels measured by RNA-seq. Data as mean ± SD, \*\**p* < 0.01, \*\*\**p* < 0.001, \*\*\*\**p* < 0.0001, \*\*\*\*\**p* < 0.00001, one-tailed Student’s *t* test.

We next made use of the HEC-1-B cell line, which expresses *α6, α12, αc1*, and *αc2*, as a single-cell model system to investigate mechanisms of gene regulation [12]. Single-cell RNA-seq analyses and maximum likelihood modeling confirmed the stochastic monoallelic expression patterns in single cells in the brain (Figure S1) [37]. We inserted CBS elements into various locations within the *Pcdhα* cluster by DNA-fragment editing and screened for single-cell CRISPR clones (Figure 1, C and D) [38]. Since inversion of *HS5-1 in situ* results in no alteration of *αc1* expression [12], and *αc2* is regulated by *HS7* in a CTCF-independent manner (Figure 1, A and B) [33], we excluded *αc1* and *αc2* from our expression analyses.

We inserted single (“F”) or tandem (“FF”) forward-oriented CBS elements into the location between the *Pcdhα* cluster and its *HS5-1* enhancer (Figure 1, E and F) and carried out quantitative high-resolution chromosome-conformation capture followed by next-generation sequencing (QHR-4C) experiments (Figure S2). QHR-4C revealed prominent long-distance chromatin interactions between *HS5-1* and the inserted CBS, and a concurrent decrease of chromatin interactions with Pcdhαtarget promoters (Figure 1, G and H and Figure S3, A-D). In addition, CBS mutations abolish these effects (Figure 1, G and H and Figure S3, A-D). Consistent with the decrease of enhancer-promoter interactions, RNA-seq revealed a significant decrease of *α6* and *α12* expression levels, and CBS mutations rescue their expression levels (Figure 1, I and J). In summary, the inserted forward-oriented CBSs block the long-distance chromatin interactions between the *HS5-1* enhancer and its target promoters, and thus function as insulators by competing with the target *Pcdhα* promoters.

We next inserted three different reverse-oriented CBS elements each into distinct locations in the *Pcdhα* cluster (Figures S3, E-I and S4). We found that each competes with the *Pcdhα* enhancer to form long-distance chromatin interactions with target promoters and thus functions as an insulator (Figures S3, E-I and S4). In addition, we inserted reverse-forward CBS pairs (“RF” or “RRFF”) into the location between the *Pcdhα* cluster and the *HS5-1* enhancer. We found that they also function as insulators (Figures S5 and S6). Finally, we inserted a pair of forward-reverse CBS elements (“FFRR”) and found that, remarkably, they still function as an insulator (Figure S7, A-D). We conclude that both forward and reverse ectopic CBSs function as insulators for the *Pcdhα* genes through CTCF-mediated directional looping. Namely, CBS insulators function in an orientation-independent manner. However, their insulation mechanisms are distinct. The forward- and reverse-oriented CBS elements form long-distance chromatin interactions with the *Pcdhα* enhancers and promoters (even passed through the oncoming convergent CBS sites, Figure S7, B and C), respectively, in an orientation-dependent manner. Thus, the location and relative orientation of the inserted CBSs determine its insulation specificity through directional looping to distinct CBS elements within the *Pcdhα* enhancers and promoters. We note that human CBS insulators are different from the *Drosophila* su(Hw) insulators in which paired insulators result in the loss of their insulation activity [2].

### Augmentation of Distal *Pcdhα* Promoter Usage by CTCF

Interestingly, similar to the *Igh* gene cluster [32], the inserted CBS elements mainly block or insulate the enhancer interactions with proximal *Pcdhα* genes (Figure 1, G and H and Figures S3F, S5B, S6B, and S7B). Surprisingly, insertion of CBS insulators augments long-distance chromatin interactions between the *HS5-1* enhancers and the distal *Pcdhα* promoters (Figure 1, G and H and Figures S3F, S5B, S6B, and S7B). To understand this puzzling phenomenon, we simulated chromosome-conformation dynamics of the *Pcdhα* cluster by a “double clamp” Cohesin dimer extrusion mechanism on a coarse-grained chromatin fragment (Figure S7E) [18–20], based on the location and relative orientation of the *Pcdhα* CBS elements that are dynamically bound by CTCF proteins (Figure 2A and Figure S8) [9, 10, 12]. Specifically, we assume that Cohesin topologically slides along the *Pcdhα* chromatin fiber until it encounters an opposite CBS element or another sliding Cohesin ring (Figure S7E) [18, 19, 39]. Remarkably, computational simulations revealed that, in addition to proximal *Pcdhα* promoter insulation, Cohesin loop extrusion results in a significant increase of chromatin interactions between the *HS5-1* enhancer and the distal *Pcdhα* promoters upon insertions of various CBS insulators (Figure 2B), consistent with the observed data from the QHR-4C experiments (Figure 1, G and H and Figures S3F, S5B, S6B, and S7B). We conclude that insertion of CBS insulators augments distal *Pcdhα* promoter-enhancer chromatin interactions.

**Figure 2.**
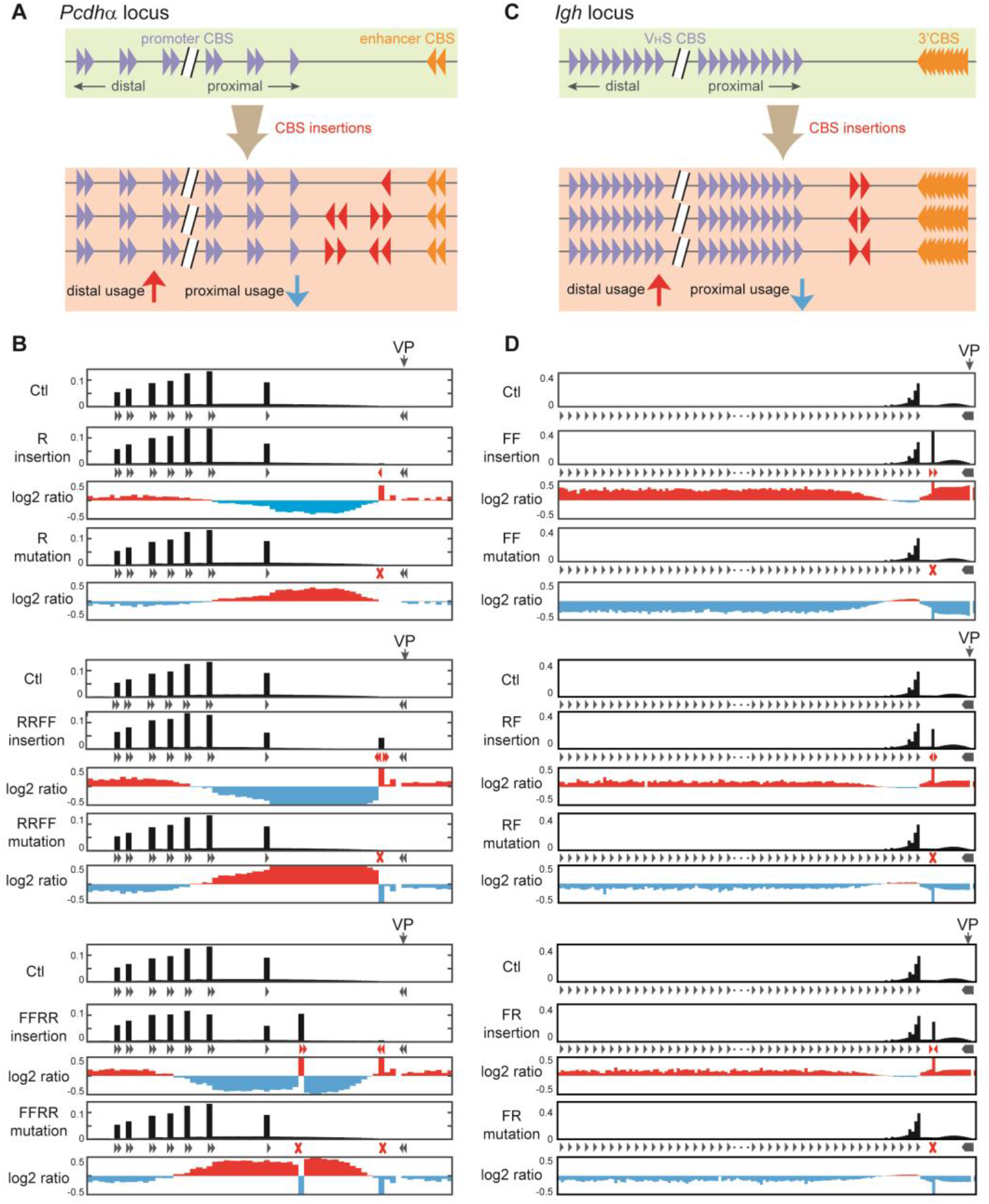
Augmentation of distal promoter usage by insertion of CTCF sites. (**A**) Schematic of increased distal and decreased proximal promoter usage upon CBS insertion in the *Pcdhα* cluster. (**B**) Computational simulation of chromatin interaction profiles using *HS5-1* as a viewpoint with insertion of different CBSs in distinct orientations and their corresponding mutations revealed CBS-dependent augmentation of distal promoter usage. (**C**) Schematic of increased distal and decreased proximal *V_H_S* usage upon insertion of various CBSs in different orientations in the *Igh* cluster. (**D**) Computational simulation of chromatin interaction profiles with 3’CBS as a viewpoint with insertion of various CBSs in different orientations and their corresponding mutations revealed similar CBS-dependent augmentation of distal promoter usage in the *Igh* cluster.

We also simulated chromosome conformation of the *Igh* cluster which also contains a large repertoire of tandem variable CBSs (Figure 2C and Figure S8) [32] and found that, similar to that in the *Pcdhα* clusters, insertion of various CBS insulators in different orientations also augments distal *V_H_* gene utilization (Figure 2D). Thus, CTCF-mediated directional looping of tandem-arrayed CBS elements determines the promoter balance of both *Pcdhα* and *Igh* clusters. We note that insertion of reverse CBSs appears to have a stronger effect on chromatin interactions between the *HS5-1* enhancer and the distal *Pcdhα* promoters than the insertion of forward ones (comparing Figure S3F to Figure 1G). Simulations of the *Pcdhα* cluster also revealed the same phenomenon (comparing Figure 2B to Figure S8, B and C). By contrast, insertion of forward CBSs appears to have a stronger effect on long-distance chromatin interactions between the 3’CBS and distal *V_H_* gene segments than insertion of reverse ones in the *Igh* cluster (Comparing the upper panel of Figure 2D to Figure S8G). The reason for this difference is unknown but may be related to different patterns of Cohesin loading by NIPBL between the *Pcdhα* and *Igh* clusters (Figure S8, A and F).

### Endogenous CTCF Sites as Protocadherin Insulators

We next tested whether each of the endogenous tandem arrays of the forward-oriented *Pcdh* CBS elements function as an insulator. We found that deletion of the *αc1* CBS element results in a significant increase of long-distance chromatin interactions between *HS5-1* and the *Pcdhα* genes upstream of *αc1* (Figure 3, A and B). In addition, this deletion results in a significant increase of *α6* and *α12* expression levels (Figure 3C). Moreover, deletion of the *α12* CBS element also results in a significant increase of chromatin interactions between *HS5-1* and the upstream *Pcdhα* genes (Figure 3, D and E) as well as of *α6* expression levels (Figure 3F). Together, these data suggest that the endogenous CBS element of *αc1* or *α12* functions as an insulator for its respective upstream *Pcdhα* genes.

**Figure 3.**
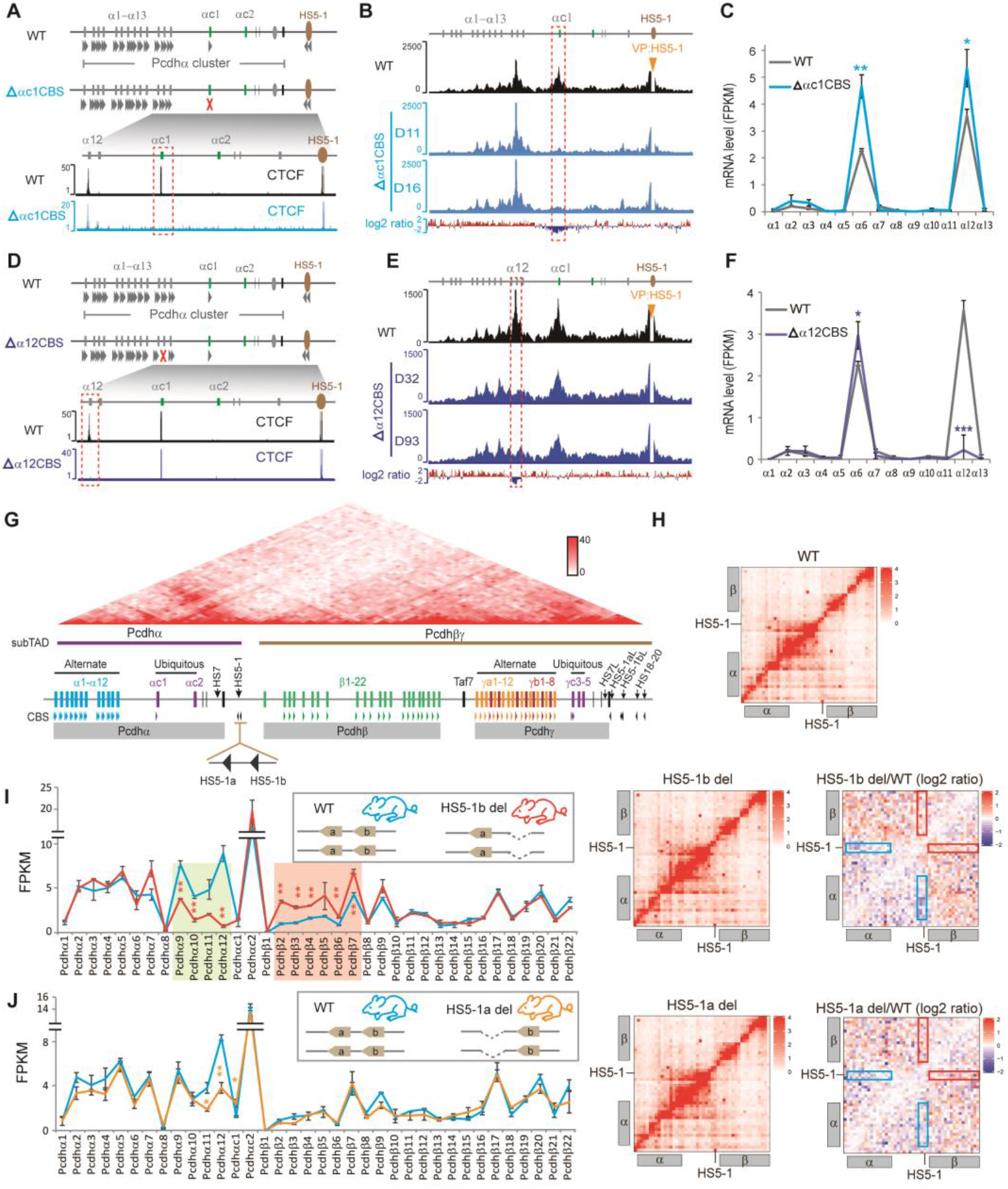
Endogenous CTCF sites as *Pcdh* insulators. (**A**) Schematic of the *Pcdhαc1* CBS deletion. The *αc1* region is highlighted. (**B**) QHR-4C with *HS5-1* as a viewpoint. Log2 ratios (deletion vs wildtype) are also shown. (**C**) RNA-seq of the wildtype (WT) and αc1-CBS deleted CRISPR cell clones. (**D-F**) Similar to (**A-C**), but with *α12* CBS deletion. (**G**) Schematic and HiC map of the three *Pcdh* gene clusters, which are organized into *α* and *βγ* subTADs, with the *HS5-1b* CBS as a boundary element. (**H**) 5C interaction profiles of the *Pcdh α* and *β* clusters. The log2 ratios of chromatin interactions of *HS5-1* with *Pcdh α* or *β* gene repertoire are highlighted by blue or red rectangles, respectively. (**I-J**) RNA-seq revealed alterations of expression levels of the *Pcdh α* and *β* gene repertoires in homozygous *HS5-1b* (**I**) or *HS5-1a* (**J**) CBS deletion mice. WT: wildtype; del: deletion. Data as mean ± SD, \**p* < 0.05, \*\**p* < 0.01, \*\*\**p* < 0.001.

To investigate whether each of the two reverse-oriented CBS elements (*HS5-1a* and *HS5-1b*) flanking the *HS5-1* enhancer also functions as an insulator, we deleted each of them in mice and performed 5C and RNA-seq experiments using mouse cortical tissues (Figure 3, G-J). Deletion of the *HS5-1b* CBS (Figure S9, A and B), which is at the boundary between the *Pcdhα* and *Pcdhβγ*subTADs (Figure 3G), results in an aberrant increase of long-distance chromatin interactions between *HS5-1* and the 5’-end isoforms of the *Pcdhβ* cluster as well as an aberrant activation of their promoters (Figure 3, H and I). By contrast, both the chromatin interactions of *HS5-1* with the proximal *Pcdhα* genes as well as their expression levels are significantly decreased (Figure 3, H and I). This suggests that the boundary *HS5-1b* CBS element is an insulator that restricts the *HS5-1* enhancer activity from the aberrant activation of the *Pcdhβ* promoters. As a control, homozygous deletion of the internal *HS5-1a* CBS element (Figure S9C) results in no alteration of the 5’-end isoforms of the *Pcdhβ* cluster (Figure 3, H and J). Therefore, although both *HS5-1a* and *HS5-1b* CBSs are required for bridging the *HS5-1* enhancer to the *Pcdhα* promoters (Figure 3, H-J), only the boundary *HS5-1b* CBS element functions as an insulator restricting the *HS5-1* enhancer activity from aberrantly activating the *Pcdhβ* genes.

### Insulators for Enhancers with no CTCF Site

We next prepared mice with a deletion of the entire *HS5-1* fragment including the two flanking CBS elements (Figure S9D). As expected, this results in a significant decrease of chromatin interactions between *HS5-1* and the *Pcdhα* genes as well as their expression levels (Figure 4, A-C). Surprisingly, we found that the expression levels of the 5’-end isoforms of the *Pcdhβ* cluster are significantly increased upon *HS5-1* CBS deletion and that the long-distance chromatin interactions between the *HS7* enhancer and the 5’-end isoforms of the *Pcdhβ* cluster are also significantly increased (Figure 4, B and C). This suggests that the two *HS5-1* CBS elements function as an insulator to restrict the activity of the *HS7* enhancer, which contains no CBS, from aberrantly activating the *Pcdhβ* promoters (Figure 4, A-C). As a control, we inverted *in situ* the entire *HS5-1* fragment including the two reverse-oriented *HS5-1* CBSs in mice (Figure 4A and Figure S9E). In contrast to the *HS5-1* deletion, neither *HS7* chromatin looping interactions with nor expression levels of the 5’-end isoforms of the *Pcdhβ* cluster are increased (Figure 4, B and D). These remarkable differences between the *HS5-1* deletion and inversion clearly show that the two endogenous *HS5-1* CBS elements block the *HS7* enhancer from the aberrant activation of the *Pcdhβ* genes and their insulation activity is orientation-independent *in vivo*, consistent with the insertions of exogenous CBSs of either orientation in cell lines *in vitro* (Figure 1 and Figures S3–S7).

**Figure 4.**
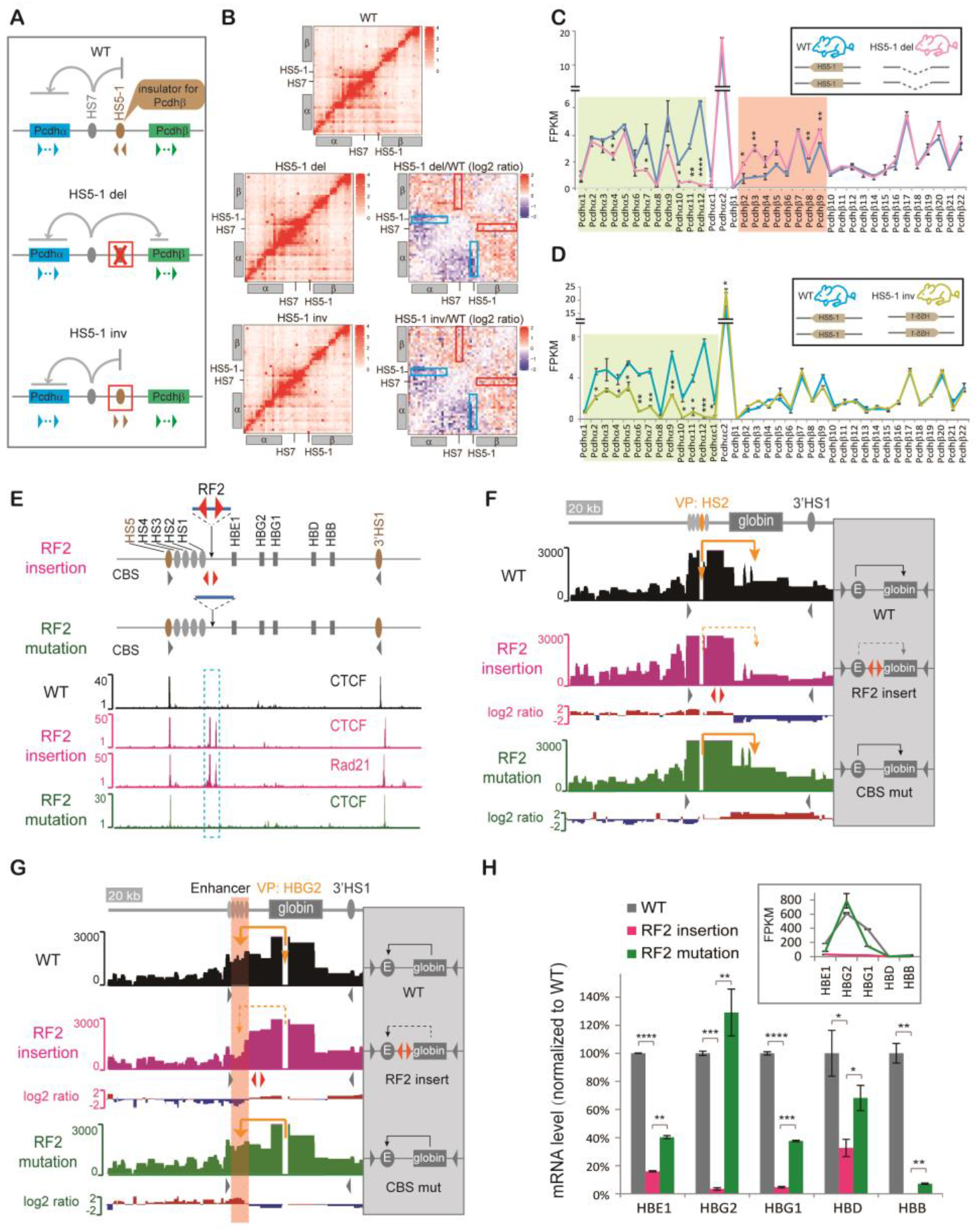
Insulators for enhancers with no CTCF site. (**A**) Schematic of the *HS5-1* CBSs as an insulator of the *HS7* enhancer for the *Pcdhβ* genes. (**B**) 5C interaction profiles of the *Pcdh a* and *β* clusters in the *HS5-1* deletion (del) or inversion (inv) mice *in vivo*. The log2 ratios of chromatin interactions of *HS5-1* with *Pcdhα* or *HS7* with *Pcdhβ* are highlighted by blue or red rectangles, respectively. (**C**) RNA-seq of cortical tissues of the WT and *HS5-1* deletion mice. (**D**) RNA-seq of cortical tissues of the WT and *HS5-1* inversion mice. (**E**) CTCF and Rad21 ChIP-seq of human single-cell *β*-globin CRISPR clones with insertion of a pair of reverse-forward CBSs (“RF2”). (**F**) The 4C-seq profiles with the *β-globin HS2* enhancer as a viewpoint. (**G**) The 4C-seq profiles with the human *β-globin HBG2* promoter as a viewpoint. (**H**) RNA-seq revealed decreased expression levels (normalized to WT) of the human *β*-globin repertoire. The actual expression levels are shown in the Inset. Data as mean ± SD, \**p* < 0.05, \*\**p* < 0.01, \*\*\**p* < 0.001, \*\*\*\**p* < 0.0001.

We next inserted a pair of reverse-forward CBSs (designated “RF2” to be distinguished from the first “RF” in Figure S5) into the location between the *HS2* enhancers, which also contains no CBS, and promoters of the *β*-globin cluster (Figure 4E). ChIP-seq confirmed the binding of CTCF/Cohesin to the inserted CBS pair but not its mutant sites (Figure 4E). QHR-4C experiments with either the *HS2* enhancer or the *HBG2* promoter as a viewpoint demonstrated a significant decrease of the *β*-globin enhancer-promoter interactions (Figure 4, F and G). Consistently, the expression levels of all *β*-globin genes are significantly decreased and the decrease is rescued by CBS mutations (Figure 4H). Finally, QHR-4C with the inserted CBS pair as a viewpoint revealed opposite chromatin looping interactions with both *5’HS5* and *3’HS1*, which contain CBSs located outside of and beyond the *β*-globin enhancer and promoter regions, respectively (Figure S10, A-C). Thus, CBS elements, if inserted between enhancers and promoters with no CBSs, also function as insulators by forming long-distance chromatin looping interactions with CBS elements located in the endogenous genome outside of and beyond respective enhancer and promoter regions. Finally, we inserted various combinations of CBSs upstream and/or downstream of the *HS7* enhancer of the *Pcdhα* cluster and found that the inserted CBSs block long-distance chromatin interactions of the *HS7* enhancer (Figure S10, D-G). In conjunction with data of the endogenous *Pcdh* CBS deletion (Figure 4, A-D), we conclude that CBS elements can function as insulators for enhancers with no CBS.

### Genome-wide CTCF Sites as Insulators

To see whether CTCF-bound CBS elements genome wide function as insulators for enhancers to ensure proper activation of promoters, we analyzed CTCF occupancy of CBSs located between neighboring enhancer-promoter pairs and their corresponding promoter activities in HEC-1-B, HEK293T, and K562 cell lines by performing ChIP-seq and RNA-seq experiments (Figure 5, A and B). Remarkably, in each of these three cell lines, promoter activities are inversely correlated with insulator scores (Figure 5B). Finally, we analyzed the published ENCODE datasets and found the same inverse correlations in six other cell lines (Figure S11) [40]. Thus, these computational analyses, in conjunction with our inversion and deletion experiments, suggest that CTCF-bound CBSs could function as insulators through directional chromatin looping across the genome.

**Figure 5.**
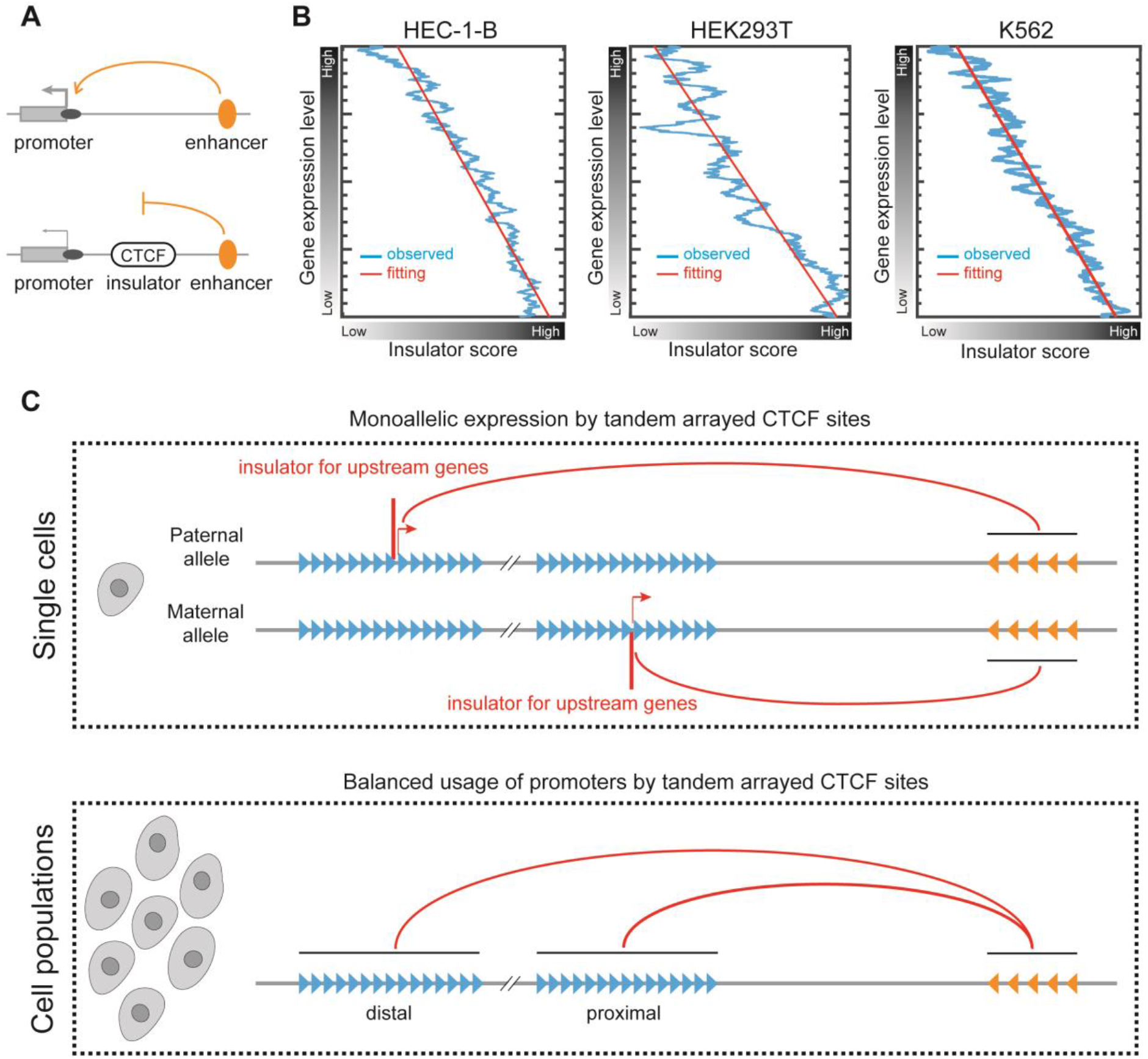
CTCF sites genome wide negatively correlate with gene expression levels. (**A**) Schematic of promoter-enhancer (P-E) pairs with or without intervening CBSs. (**B**) Negative correlations between insulator scores and gene expression levels in each of the HEC-1-B, HEK293T, and K562 cell lines. (**C**) A model of CTCF directional looping insulation mechanism. Directional CTCF looping underlies stochastic monoallelic *Pcdh α* and *βγ*gene expression in single cells and balanced variable promoter usage in cell populations in the brain, as well as balanced variable gene segment usage of the *Igh* cluster in primary pro-B lymphocyte populations. In particular, at the single-cell level, stochastic and monoallelic CTCF-mediated directional chromatin looping underlies activation of one and only one variable promoter in each chromosome [34, 36]. At the cell-population level, large repertoires of tandem-arrayed CBSs ensure balanced variable promoter usage of the *Pcdh* and *Igh* clusters in the nervous and immune systems, respectively.

In summary, by a combination of CRISPR DNA-fragment editing and chromosome-conformation capture experiments, we showed that ectopic and endogenous CBS elements function as insulators in an orientation-independent manner through CTCF-mediated directional chromatin looping. In conjunction with computational simulation of “double clamp” loop extrusion, we showed that tandem-arrayed variable CBSs ensure accessibility of repertoires of tandem-arrayed promoters and their balanced usage.

## DISCUSSION

Considerable progress has been made in understanding the regulatory mechanisms of clustered stochastic monoallelic *Pcdh* gene expression in single neurons in the brain [12, 27, 34, 36, 41]. The allelic insulation by CTCF-mediated directional looping may be epigenetically regulated by methylation of CBS elements [34]. Some CBS elements, such as the *Pcdh* enhancer *HS5-1b* site, contain no CpG dinucleotides, and thus cannot be regulated by CpG methylation [34]. Consequently, it has constitutive and cell-invariant CTCF/Cohesin occupancy and functions as an insulator for the downstream *Pcdhβ* genes (Figure 3, H and I). Other CBSs are regulated by DNA methylation, and have a cell-specific pattern of the CTCF/Cohesin occupancy in single neurons [27]. For example, each CBS flanking all alternate variable promoters of the *Pcdhα* cluster contains a CpG dinucleotide that is methylated in specific subpopulations of neurons in the brain [27, 34, 42]. Therefore, the upstream boundary CBS element of each *Pcdhα* loop domain is cell-specific and distinct for single neurons, thus functioning as an insulator for its respective upstream genes. Consistently, our computational analysis based on stochastic looping in each chromosomal allele and the expression of one or two isoforms in single cells in the mouse brain suggest that the alternate members of the *Pcdhα* family are expressed monoallelically in individual neurons (Figure S1) [37]. This explains the long-standing puzzle of stochastic monoallelic expression of alternate members of the *Pcdhα* family in individual neurons in the brain [36]. Specifically, for any unmethylated promoter CBS, it forms long-distance chromatin interactions with downstream enhancers and is activated through chromatin looping. These interactions function as an insulator for all of its upstream promoters in each chromosomal allele in single cells while all other variable promoters are insulated and silenced. Therefore, only one isoform could be expressed from each chromosomal allele in individual neurons (Figure 5C).

Overwhelming evidence suggests that the function of insulators is orientation-independent but the chromatin looping of CBS elements is directional [7, 11, 12, 14, 23, 32, 43]. CTCF mediates specific directional loop formation through asymmetric anchoring of the ring-shaped Cohesin complex, which topologically slides along chromatin fibers to extrude loops [3, 12, 18, 19, 32, 44]. Our data are consistent with the observed predominant chromatin interactions between forward-reverse CBS pairs [11, 12]. In addition, there are numerous cases of tandem CBS clusters [12, 16, 21, 32]. Since the binding of CTCF to genome-wide CBSs is not static but rather dynamic [9, 10] and there are variable permeabilities of CTCF extrusion barriers [20], this suggests that Cohesin may slide through the proximal CBS in tandem CBS clusters. Curiously, our computational simulation and chromosome-capture experiments revealed that tandem-arrayed CBSs ensure balanced usage of their associated promoters in cell populations (Figure 5C). In addition, deletion of CBS elements in the *Igh* cluster [32] leads to decreased distal and increased proximal promoter usage. Thus, directional looping between convergent CBSs underlies insulator function and tandem-arrayed CBSs ensure balanced promoter accessibility.

## MATERIALS AND METHODS

### Cell culture

Human endometrial HEC-1-B cells (ATCC) were cultured in MEM medium (Hyclone), supplemented with 10% (v/v) FBS (Gibco), 2 mM glutamine (Gibco), 1 mM sodium pyruvate (Sigma), and 1× penicillin-streptomycin (Gibco). Human K562 and HEK293T cells (ATCC) were cultured in DMEM medium (Hyclone) supplemented with 10% (v/v) FBS and 1 × penicillin-streptomycin. Cells were maintained at 37 °C in a humidified incubator containing 5% (v/v) of CO2, and were passaged every three days.

### *In vitro* transcription of sgRNA pairs and Cas9 mRNA for microinjection

The preparation of sgRNA pairs and Cas9 mRNA were recently described [38]. Briefly, to obtain sgRNAs for microinjection of zygotes, we performed *in vitro* transcription using DNA templates generated by PCR with a forward primer containing a T7 promoter followed by targeting sequences and a common reverse primer. *In vitro* transcription was performed with the MEGA shortscript Kit (Life Technologies) using T7 polymerase incubated at 37 °C for 5 hr. The template DNA was removed by digestion with DNaseI. The transcribed sgRNAs were purified with the MEGAclear Kit (Life Technologies) and eluted in TE bumffer (0.2 mM EDTA).

To obtain Cas9 mRNA for microinjection of zygotes, the Cas9 coding sequence was clone into pcDNA3.1 plasmid under the control of the T7 promoter. The plasmid was then linearized by XbaI and used for *in vitro* transcription with the mRNA transcription system according to the manufacturer’s instructions (Life Technologies). After digestion of DNA template, the transcribed Cas9 mRNA was purified with the MEGAclear Kit (Life Technologies).

### Generation of the CBS deletion and inversion mice by CRISPR DNA-fragment editing

Mice were maintained at 23 °C in a 12-hour (hr) (7:00-19:00) light and 12-hr (19:00-7:00) dark schedule in an SPF mouse facility. For each CRISPR deletion or inversion of CBS elements, Cas9 mRNA (100 ng/μl) and a pair of sgRNAs (50 ng/μl each) targeting the region flanking the CBS elements were injected into the cytoplasm of one-cell embryos of the C57BL/6 mice. After recovering for 2 hr at 37 °C incubator, the embryos were then implanted into the oviducts of the pseudo-pregnant ICR mice. The new born F0 mice were then screened for targeted deletions or inversions by PCR using specific primer pairs. The amplified PCR products were then cloned and confirmed by Sanger sequencing. The F0 mice with targeted deletions or inversions were maintained and crossed to obtain F1 mice. F1 mice were genotyped again for heterozygous deletion or inversion. Heterozygous F1 mice were then crossed to obtain homozygous F2 mice. For all of the 5C and RNA-seq experiments, only wildtype littermates were used as controls. All animal experiments were approved by the Institutional Animal Care and Use Committee (IACUC) of Shanghai Jiao Tong University (protocol#: 1602029).

### Single-cell RNA-seq

Single-cell RNA-seq experiments were performed as previously described [37]. Briefly, for neurons, P0 mouse brain was dissected, and the tissue from cerebral cortex was digested with 0.013% of collagenase in Neurobasal Medium (Gibco) at 37°C for 3 min. The collagenase was neutralized by adding excess amount of Neurobasal Medium. Single-cell suspension was made by gentle pipetting, and then filtered through 100-μm cell strainers (BD Biosciences). For HEC-1-B cells, trypsin was added to the culture dish and the single cells were suspended in the culture medium. Single cells were then picked under the microscope by using a micro capillary pipette into the thin-walled PCR tube containing 2 μl of cell lysis buffer, 1 μl of oligo-dT primer and 1 μl of dNTP mix. After reverse transcription, the cDNA was pre-amplification by PCR. The cDNA library was then purified, tagmented, and ligated with adapters using the Nextera XT DNA Library Preparation kit (Illumina FC-131-1096). Finally, the adapter-ligated fragments were further amplified by PCR and purified with AMPure XP beads (Beckman). The single-cell RNA-seq libraries were pooled and sequencing using an Illumina Hiseq 2500 platform.

### Plasmid construction

The plasmids of sgRNAs for cell transfection experiments were constructed as previously described [38, 45]. Briefly, pairs of complementary oligonucleotides for generating sgRNAs were annealed with 5’ overhangs of ‘ACCG’ and ‘AAAC’, and cloned into a BsaI-linearized pGL3 vector under the control of the U6 promoter. To insert CBS elements into distinct genomic regions, circular donor plasmids with about 2 kb homologous arms flanking the inserted sequence were used as donors for CRISPR-based homologous recombination. To construct donor plasmids, we amplified the CBS elements, as well as the genomic sequences flanking the insertion site by PCR. CBS elements and the two homologous arms with 20 bp of overlapping sequences were jointed together with the EcoRI and HindIII digested Puc19 vector using the multi-fragment recombination system (Vazyme). All of the plasmids constructed were confirmed by Sanger sequencing.

### Screening CBS insertion and deletion single-cell clones by CRISPR DNA-fragment editing

Generation of the CRISPR single-cell clones with CBS element insertions and deletions was performed as previously described [12, 38]. Briefly, cells were transfected with plasmid mix using Lipofectamine 3000 reagents (Thermo) in a 12-well plate. For CBS insertions and mutations, Cas9 (0.3 μg) and donor plasmids (0.5 μg) were co-transfected with one sgRNA construct (0.2 μg) targeting the insertion site. For CBS deletions, Cas9 plasmids (0.4 μg) were co-transfected with two sgRNA constructs (0.3 μg each) targeting the two ends of the deletion fragments. The sgRNA constructs contained puromycin-resistant gene which can be used for selection. 48 hr after transfection, puromycin (Sigma) was added to the culture medium at a final concentration of 2 μg/ml. The culture medium was replaced every day with puromycin for a total of 3 days. Puromycin was then removed and cells were cultured in normal culture medium for 2 days. The cells were then suspended into single-cell solution and plated into 96-well plates at the concentration of about one cell per well. Two weeks after plating, single-cell clones were marked manually under the microscope, and replaced with fresh culture medium. Four weeks after plating, the single-cell clones were screened for insertion, mutation, or deletion by PCR. At least two individual clones for each insertion, mutation, or deletion were obtained and analyzed. We screened for a total 1,948 single-cell clones, and 80 homozygous clones were obtained and analyzed.

### ChIP-seq experiments

ChIP experiments were performed as previously described [34] with modifications. Briefly, 4 × 10^6^ of cells were cross-linked by 1% formaldehyde in 10% FBS/PBS for 10 min at room temperature. Cells were then lysed twice with ice-cold lysis buffer (20 mM Tris-HCl, 2 mM EDTA, 1% Triton X-100, 0.1% SDS, 0.1% sodium deoxycholate and 1× protease inhibitors, pH 7.5) for 10 min with slow rotations. The lysed cells were then sonicated to obtain DNA fragments of about 200-500 bp using the Biorupter system (high energy, with working time of 30 seconds and resting time of 30 seconds, 30 cycles). After removal of the insoluble debris, the lysate was incubated with specific antibodies against CTCF (07-729; Millipore), RAD21 (ab992; Abcam), or H3K27ac (ab4729; Abcam) and purified by protein A-agarose beads (16-157; Millipore). NIPBL ChIP-seq were from recently published data [46]. ChIP DNA was extracted and prepared for high throughput sequencing using a DNA library preparation kit for Illumina (NEB). ChIP-seq libraries were sequenced on a HiSeq X Ten Platform (Illumina).

### Quantitative High-resolution Chromatin Conformation Capture Copy (QHR-4C)

The QHR-4C protocol was modified from previously published method [32]. Briefly, one hundred thousand cells from various CRISPR single-cell clones were centrifuged at 500 g for 5 min and the pellets were used for QHR-4C experiments. The cell pellets were suspended for crosslinking in 900 μl 2% formaldehyde at room temperature for 10 min. The crosslinking reaction was stopped by adding and mixing with 100 μl of 2M glycine for a final concentration of 200 mM. The fixed cells were spin down at 800 g at 4 °C for 5 min and washed twice by suspending briefly in 1 ml ice-cold PBS. Cells were then permeabilized twice with 200 μl ice-cold 4C permeabilization buffer each for 10 min (50 mM Tris–HCl pH 7.5, 150 mM NaCl, 5 mM EDTA, 0.5% NP-40, 1% Triton X-100, and 1× protease inhibitors). The cells were then digested overnight at 37 °C with DpnII while shaking at 900 rpm. After inactivation of DpnII at 65 °C for 20 min, proximity ligation was carried out for 24 hr at 16 °C with T4 DNA ligase in 0.1 ml 1 × T4 ligation buffer. The ligated product was then reverse cross-linked and DNA was extracted using phenol-chloroform. Glycogen was added to facilitate DNA precipitation. The precipitated DNA was then dissolved in 50 μl water, and sonicated using the Biorupter system (with low energy setting at a train of 30-second sonication with 30 second intervals for 12 cycles) to obtain DNA fragments ranging from 200 to 600 bp.

The captured fragments were linear-amplified using a 5’ biotin-tagged primer complementary to the viewpoint fragment in 100 μl of PCR system for a total of 60 cycles. This primer should be neither too close to the DpnII site to facilitate the nested PCR at the final amplification step, nor too far away from the DpnII site to maximize the product amount. The amplification products were denatured by incubating at 95 °C for 5 min, and immediately chilled on ice to obtain ssDNA. The ssDNA was then enriched and purified with Streptavidin Magnetic Beads (Invitrogen) according to the manufacturer’s instructions.

The ssDNA on-beads was then ligated in 15 μl with 0.1 μM of adapters at 16 °C for 24 hr. The adapters were generated by annealing two complementary primers matching Illumina P7 sequences. After ligation, free adapters were removed by washing the beads twice with the B/W Buffer (5 mM Tris-HCl, 1 M NaCl, 0.5 mM EDTA, pH 7.5). Finally, the QHR-4C libraries were generated by one-step PCR amplification (94 °C, 2 min; 94 °C, 10 sec, 60 °C, 15 sec, 72 °C, 1 min for 19 cycles; and a final extension at 72 °C, 5 min) with captured ssDNA on beads as the template and a pair of PCR primers. The forward primer matches the Illumina P5 and the viewpoint sequence adjacent to the viewpoint DpnII site with barcodes, and the reverse primer matches Illumina P7 with indexes. The PCR products were purified with a PCR purification kit (Qiagen). About 100 QHR-4C libraries with different combinations of barcodes and indexes were pooled and sequenced on an Illumina HiSeq X Ten Platform. All of the QHR-4C experiments for each CRISPR clone were performed with biological replicates.

### Circularized Chromosome Conformation Capture (4C)

The 4C experiments were performed as previously described [12, 14]. Briefly, cells were counted and about 2×10^6^ cells were used for each 4C experiment. After cross-linking with 2% formaldehyde, cells were lysed twice with cold lysis buffer, digested with DpnII, ligated with T4 DNA ligase. The ligated samples were purified using the High-Pure PCR Product Purification kit (Roche). The 4C-seq libraries were generated by PCR using a high-fidelity DNA polymerase (Vazyme). All of the 4C experiments were performed with biological replicates. 4C-seq libraries were sequenced on the HiSeq X Ten Platform.

### Chromosome Conformation Capture Carbon Copy (5C)

5C experiments were performed as previously described [47]. Briefly, a total 46 forward and 46 reverse primers covering the mouse *Pcdh α* and *β* clusters were designed by My5C tools (http://my5c.umassmed.edu). All forward primers contain CGG and a T7 Universal primer sequence (CGGTA ATACG ACTCA CTATA GCC) at its 5’-end followed by a unique sequence. All reverse primers contain CTT at 5’-end followed by a unique sequence and a complementary T3 Universal sequence (TCCCT TTAGT GAGGG TTAAT A). All primers were 5’-phosphorylated.

The P0 mouse cortex tissues were dissociated to obtain single-cell suspension as previously described in the single-cell RNA-seq experiments. A total of 10^7^ cells were cross-linked and digested with HindIII (NEB). After inactivating HindIII, the digested DNA was ligated with T4 DNA ligase and purified. As a control, DNA of six bacterial artificial chromosomes (BACs) covering the *Pcdh* clusters was also digested, ligated, and purified. The purified mouse cortical DNA was mixed with 1 μg of salmon sperm DNA (Sigma). The control BAC DNA (5 ng) was mixed with 1.5 μg of salmon sperm DNA. These samples were then each mixed with 1.7 fmol of each 5C primer and 1 μl of 10×5C annealing buffer (20 mM Tris-acetate pH7.9, 50 mM potassium acetate, 10 mM magnesium acetate, 1 mM DTT) in a total volume of 10 μl and denatured at 95 °C for 5 min. Annealing was performed by incubation at 48 °C for 16 hr. The annealed DNA was ligated by adding Taq DNA ligase (NEB) in the 5C ligation buffer (25 mM Tris-HCL pH7.6, 31.25 mM potassium acetate, 12.5 mM magnesium acetate, 1.25 mM NAD, 12.5 mM DTT and 0.125% Triton X-100). The ligation reaction was performed for 1 hr at 48 °C followed by incubation for 10 min at 65 °C to stop the ligation reaction. The ligated products were amplified by PCR with Illumina primer pairs and purified with PCR purification kit (QIAGEN). 5C libraries were sequenced with the 90 bp pair-end mode by Hi-seq 2500 platform of Illumina. All 5C experiments were performed with biological replicates.

### *In situ* Hi-C

Hi-C experiments were performed as previously described [12]. Briefly, cells were crosslinked with 1% formaldehyde for 10 min at RT. After permeabilizing the nuclei, DNA was digested with MboI (NEB). DNA fragments were labeled with biotin and ligated *in situ*. After reversing crosslinks, DNA was purified and sonicated. Biotinylated ligation junctions were enriched with streptavidin beads on a magnetic stand. DNA on beads was then ligated with adapters and amplified with PCR for Illumina sequencing. Normalized contact matrices were constructed after removing intrinsic biases in Hi-C data. Topologically associated domains (TADs) and subTADs were called as described [11].

### RNA-seq experiments

RNA-seq experiments were performed as previously described [12] with modifications. Briefly, total RNA from mouse cortex or culture cells was extracted using Trizol reagents (Life Technologies) following the manufacturer’s instructions. Total mRNA was prepared from 1 μg total RNA using poly(A) mRNA magnetic isolation reagents (NEB) and fragmented at 94 °C for 15 min. RNA was then reverse-transcribed into cDNA with random primers. After end repairing and A-tailing, cDNA was ligated with adapters and amplified by PCR with Illumina sequencing primers. All RNA-seq experiments were performed with biological replicates. RNA-seq libraries were sequenced on a HiSeq X Ten Platform.

### High-throughput Sequencing and Data Analyses

High-throughput analyzed pipelines were the same as previously described [12, 34] with some modifications. Briefly, reads that passed the Illumina quality filter were considered for alignments. For 4C-seq data, reads were aligned to the reference human genome (GRCh37/hg19) using the Bowtie program (version 1). The reads per million (RPM) value was calculated using the r3Cseq program (version 1.20) in the R package (version 3.3.3). For QHR-4C data, duplicated paired-end reads were removed by FastUniq (version 1.1) program, and only the unique reads were used for analyses using the Bowtie and r3Cseq program. For 5C data, the reads matched every primer-pair were counted and the total matched reads of each sample were adjusted to 1 million. The counting results were then normalized to the BAC control. The heatmap of 5C results and log2 ratios were drawn with the R package ggplot2. For ChIP-seq data analyses, reads were mapped to the reference genome (GRCh37/hg19) or the modified genome with insertions using the Bowtie program. Peaks were called by the MACS program (version 1.4.2) with a cutoff *P* value of 10^−5^. For RNA-seq and single-cell RNA-seq data, reads were aligned using Hisat2 (version 2.0.4) to the human genome (GRCh38/hg38) or mouse genome (GRCm38/mm10), and the FPKM value was calculated using the Cufflinks program (version2.1.1).

### Maximum likelihood modeling of stochastic expression

Since single-cell RNA-seq data of each neuron are resulted from the combined expression of two sets of paternal and maternal chromosomes, upon the assumption that two chromosomal sets express independently, we first decomposed the RNA-seq data of single cells from the anterior lateral motor and primary visual cortices [37] into the expression of each chromosomal sets.

Let *G* be the total number of considered genes (for example, *G* = 12 in the mouse *Pcdhα* cluster). Define the whole gene set as 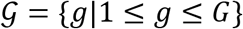. Because there are 81.56% and 86.64% single cells from anterior lateral motor and primary visual cortices express no more than 2 *Pcdhα* isoforms (Figure S1), respectively, we assume that the *Pcdhα* cluster on a single chromosomal allele expresses at most *H* genes (*H* = 2 here in the *Pcdhα* cluster). Define 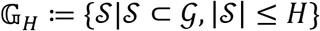 where 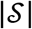 is the total number of elements in the set 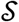, as the set of all the subsets of 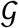 that contain less than *H* elements. In other words, 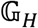 gives all possible gene sets that can be expressed from a single chromosomal allele.

Define the Cartesian product 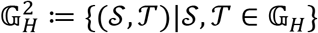 as all possible combinatorial expression sets from both chromosomal alleles. For 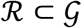, let *N_ℛ_* be the number of single cells that express the gene set *ℛ*. Define 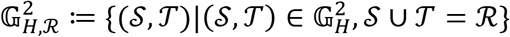. 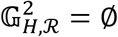 if and only if 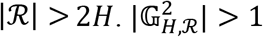 means that there are more than one way of the *Pcdhα* isoforms to be expressed from both chromosomal alleles to achieve the total expressed gene set *ℛ*. Define 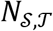 as the number of single cells that the first chromosomal allele expresses gene set 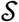 and the second chromosomal allele expresses gene set 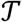. 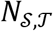 is hidden. By definition, 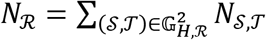. Define 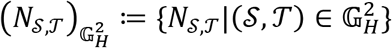 and 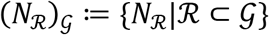 under the independent assumption 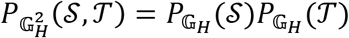, where 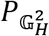 and 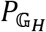 are distributions (probability measures) on 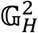 and 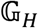. We choose 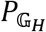 which maximizes the likelihood 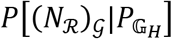. This is achieved by alternately maximizing the complete likelihood 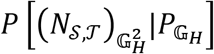 over 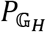, and calculating the conditional expectation 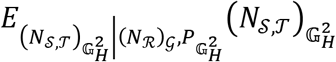 [48]. To be exact, do 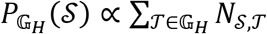 and 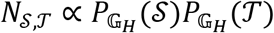 until convergence. Note that the second line is done under the constraint 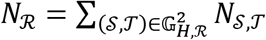. We initially assume that 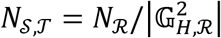 for 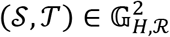.

### Computational polymer simulation of tandem arrayed CTCF sites

We developed a method to simulate long-distance chromatin interactions based on Cohesin loop extrusion on a coarse-grained DNA fragment. Our method focuses on CTCF/Cohesin distribution on one-dimensional DNA fiber and does not involve any physical property of the DNA polymer in a 3D space. The modeled DNA fragment is divided into roughly equal bins. Long-distance chromatin interactions between one bin and all other bins are determined by a graph-based method according to simulated Cohesin distribution across the DNA polymer.

### “Double clamp” Cohesin extrusion

Because two Cohesin rings can interact with each other to form a dimer [49], we proposed that, two tethered Cohesin rings, each topologically embracing a double-stranded DNA (dsDNA) within its lumen (Figure S7E), are loaded stochastically on any bin or by NIPBL and start to extrude DNA in opposite directions. The extrusion process is continuous until blocked by oriented CBS which bound CTCF protein in an antiparallel manner [10]. This model is based on the following observations:

First, Cohesin core subunits Smc1, Smc3, and Rad21 interact with themselves in an SA1/SA2-dependent manner [49]. Second, Cohesin ring can be loaded onto dsDNA in a topological as well as non-topological manner [39]. Third, closely-related Condensin mediates loop extrusion asymmetrically, with one side anchored onto dsDNA and the other side reeled in dsDNA [50]. Fourth, most Cohesin complexes are loaded at sites not overlapped with CTCF peaks [46]. Finally, loop extrusion requires ATP and only the head domain of Cohesin has the ATPase activity [46]. Thus, the “double clamp” loop extrusion suggests that a pair of tethered Cohesin rings can actively reel dsDNA in from both sides until blocked and anchored by CTCF.

### A pair of Cohesin rings extrudes dsDNA on a coarse-grained chromatin fiber

We simulated long-distance chromatin interaction profiles between the *HS5-1* enhancer and variable promoters of the *Pcdhα* cluster by coarse-grained modeling. The entire *Pcdhα* region was divided roughly into 64 bins, each of which spans about 5 kb, such that each *Pcdhα* promoter and its associated CBSs were assigned into an individual bin. We also simulated the *Igh* cluster by dividing its DNA region into 120 bins. Two tethered rings of Cohesin dimer independently extrude a DNA loop in opposite directions in an ATP-dependent manner until blocked by CTCF or dropped off from the coarse-grained chromatin fiber (Figure S7E) [19, 51]. In addition, Cohesin rings cannot pass through each other during extrusion, but are allowed to occupy the same bin. Finally, we assume that only CTCF-bound CBS can block topological Cohesin sliding in an orientation-dependent manner [11, 18, 19].

### Calculating permeability of Cohesin ring sliding through oriented CTCF array

It was recently reported that CTCF binding to dsDNA is much more dynamic than Cohesin and that the residence time of Cohesin on DNA fiber is at least 10 fold more than CTCF [9]. The dynamic binding of CTCF to oriented CBS provides hindrance for Cohesin sliding. This is generally described by the so-called permeability, *i.e*. the probability that CBS permits Cohesin ring to pass through. If there are *n* consecutive CBSs *c*_1_, *c*_2_, ⋯, *c_n_*, from distal to proximal, with permeability of *p*_1_, *p*_2_, ⋯, *p_n_*, respectively, we want to know the mean attempting times *x_n_* for Cohesin ring to slide through the entire CBS array from proximal to distal. For the first attempt, the proximal CBS has a probability *p_n_* to allow Cohesin ring to pass through. Thus, the Cohesin needs *x*_*n*−1_ attempting times in average to slide through the remaining CBS array *c*_1_, *c*_2_, ⋯, *c*_*n*−1_. Otherwise, the proximal CBS *c_n_* has the probability (1 − *p_n_*) to block Cohesin ring passing through. Thus, one attempting time has been used and Cohesin still needs *x_n_* attempting times in average to slide through the entire CBS array *c*_1_, *c*_2_, ⋯, *c_n_*. In summary,

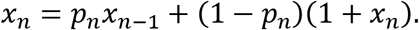

Since *x*_0_ = 1, by mathematical induction, 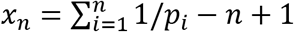. Then one obtains the overall permeability 1/x_n_ of CBSs *c*_1_, *c*_2_, ⋯, *c_n_*.

Let *k* be the sliding rate of Cohesin ring through a bin with no CBS, then the sliding rate at a bin with *n* CBSs, whose permeabilities are *p*_1_, *p*_2_, ⋯, *p_n_*, is approximately *k*/*x_n_*.

### Inferring transition rate constants of Cohesin

In order to simulate long-distance chromatin contact dynamics, we need to estimate the rate constants of Cohesin loading, sliding, and dropping. The real time scale is not necessary for our simulation. Without loss of generality, we always set the Cohesin dropping rate *d* = 1.

The concepts of Cohesin separation and processivity are introduced to characterize loop extrusion [19, 20]. Accordingly, Cohesin separation *S* is the mean distance between consecutive sliding Cohesin complexes on a chromatin fiber and Cohesin processivity *λ* is the mean size of the extruded loops.

Let *I* be the total number of bins in the coarse-grained modeling region, and all bins are numbered from upstream to downstream. Since a pair of tethered Cohesin rings extrudes DNA fiber, let 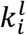 for *1* ≤ *i* ≤ *I* be the CBS-free sliding rate of the left Cohesin ring from bin *i* to *i* − 1, where bin 0 is located immediately upstream of the first bin of the region. Denote the size of bin *i* by *b_i_* for 1 ≤ *i* ≤ *I*. Let the size of bin 0 *b*_0_ be equal to the average size of all bins in the modeled region, namely, 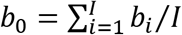. Assume that Cohesin always slides between the centers of consecutive bins, then the distance from bin *i* to bin *i* − 1 is (*b*_*i*−1_ + *b_i_*)/2. Thus, 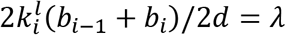, and we have the CBS-free sliding rate of the left Cohesin ring 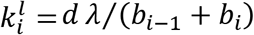.

Similarly, we have the CBS-free sliding rate of the right Cohesin ring 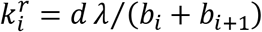 from bin *i* to *i* + 1, where bin *I* + 1 is located immediately downstream of the last bin of the region. We have 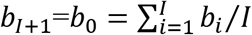.

Let 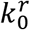 and 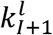 be the rates of Cohesin ring sliding into the modeled region from the left and right sides, respectively. In practice, we usually let 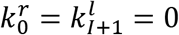 for simplicity.

Assuming uniform Cohesin loading, the loading rate *a_i_* of Cohesin to adjacent bins (*i, i* + 1) for *i* ∈ [0, *I*] depends on the separation and bin sizes by *a_i_* = *d*(*b_i_* + *b*_*i*+1_)/2*S*.

### Estimation of non-uniform Cohesin loading

In the above section, we have calculated the uniform Cohesin loading rate to adjacent bins based on the assumption that Cohesin loading is uniform across the entire modeled region. However, this is generally not true because certain protein factors such as NIPBL may promote Cohesin loading.

Suppose there are *G* types of protein factors influencing Cohesin loading. Let *γ_g_* < 1 be the ratio of Cohesin loading determined by type g for *g* ∈ [1, *G*]. We first calculate the uniform Cohesin loading rate *a_i_* for *i* ∈ [0, *I*] as in the above section. Next, we let 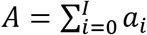 be the total loading rate for a given separation. Thus, *γ_g_A* is the part determined by type *g*. Denote the enrichment of type *g* protein factor at bin *i* by 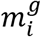 for *g* ∈ [1, *G*] and *i* ∈ [1, *I*]. Given that Cohesin recruited by type *g* at bin *i* may either bind adjacent bins (*i* − 1, *i*) or (*i, i* + 1), we define for *i* ∈ [0, *I*] that

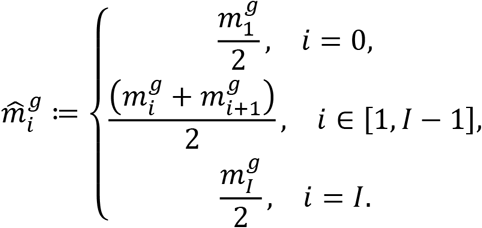

Normalize 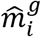 by 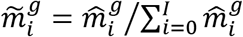. Then the binding rate *a_i_* is corrected by

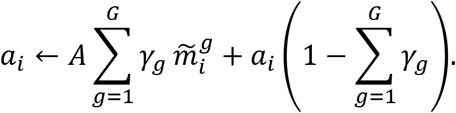

### System description and updating

Let *M_i,j_* for 1 ≤ *i* ≤ *j* ≤ *I* be the number of Cohesin ring pairs connecting bins *i* and *j*. In addition, let *L_i_* or *R_i_* for 1 ≤ *i* ≤ *I* be the number of Cohesin ring pairs connecting bin *i* and a bin downstream or upstream of the modeled region. The state of the system is determined by *M_i,j_, L_i_, R_i_*. The system may change in three ways: loading, sliding, and dropping of Cohesin.

We first describe the sliding of Cohesin ring. For *i* ∈ [1, *I*], define 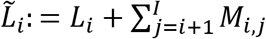 and 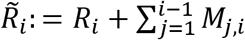 as the total number of the left and right rings at bin *i*. A left ring at bin *i* may slide to bin *i* − 1 with the rate 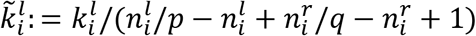 if and only if 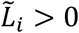 and 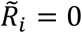, where 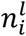 and 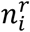 are the quantities of forward and reverse CBSs at bin *i*, and *p* and *q* are the permeabilities of opposite and non-opposite CBSs. Sliding of the left Cohesin ring from bin *i* to *i* − 1 will update *M_i,j_, L_i_, R_i_* as following. If *L_i_* > 0, then among all Cohesin ring pairs with the left ring at bin *i*, the most outside one has the right ring located downstream of the modeled region. Thereby *L_i_* decreases by 1, and if *i* > 1, increases by 1. If *L_i_* = 0, then the downstream ring of the most outside Cohesin is at bin 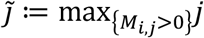. Thereby 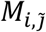 decreases by 1, and if *i* > 1, 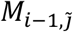 increases by 1; otherwise, 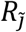 increases by 1. The sliding of the downstream ring with rate 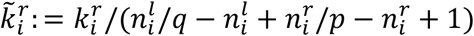 can be described similarly. Single Cohesin rings sliding into the modeled region from outside are neglected 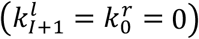 for simplicity.

Next we consider Cohesin loading. The two Cohesin rings may bind to adjacent bins (*i, i* + 1) for *i* ∈ [1, *I* − 1] with rate *a_i_*, which increases *M*_*i,i*+1_ by 1. Cohesin may bind to adjacent bins (0,1) and (*I, I* + 1) with rates *a*_0_ and *a_I_*, which respectively increase *R*_1_ and *L_I_* by 1.

Finally, all Cohesin rings may drop off with rate *d*. If a pair of Cohesin rings connecting bins *i* and *j* for 1 ≤ *i* ≤ *j* ≤ *I* drops off, then *M_i,j_* decreases by 1. If a pair of Cohesin rings connecting bin *i* and a bin downstream or upstream to the modeled region, drops off, then *L_i_* or *R_i_* decreases by 1.

### Distances between bins

The distance between adjacent bins *i* and *i* + 1 is defined as *D*_*i,i*+1_: = (*b_i_* + *b*_*i*+1_)/2. The distance between nonadjacent bins *i* and *j* connected by a pair of Cohesin rings is defined as *D_i,j_*: = *C*. If two adjacent bins *i* and *i* + 1 are connected by Cohesin, then their distance is defined as *D*_*i,i*_+1: = min[(*b_i_* + *b*_*i*+1_)/2, *C*].

For two bins *i* and *j* which are neither adjacent nor connected by Cohesin, their distance is defined as the length of the shortest path between them. If *i*_0_ = *i*, *i*_Θ_ = *j*, and ∀*θ* ∈ [0, Θ − 1], bins *i_θ_* and *i*_*θ*+1_ are adjacent or connected by Cohesin, then bins *i*_0_, *i*_1_, *i*_2_, ⋯, *i*_Θ_ are defined as a path between bins *i* and *j*. Therefore, the distance *D*_*i_θ_, i*_*θ*+1__ between bins *i_θ_* and *i*_*θ*+1_ has been defined ∀*θ* ∈ [0, Θ − 1], and the length of the path is defined as 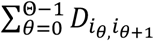. Fixing bin *i*_0_ as a viewpoint, the shortest path from the viewpoint *i*_0_ to all other bins can be efficiently searched by Dijkstra’s algorithm from the graph theory.

### Visualization of long-distance chromatin interacting profiles with a viewpoint

To visualize simulated chromatin interactions with a viewpoint, we cumulate the time *T_i_* during which bin *i* is close to the viewpoint *i*_0_ after the system enters the steady state. Because the system is ergodic, the long-term average of single-cell simulation is the same as the average of cell populations, the situation that we performed QHR-4C experiments on cell populations. We first set *T_i_* = 0 and initialize the system state *M_i,j_, L_i_, R_i_*. To allow the system to enter the steady state, we skip the first 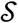 iterations. We record every 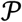 iterations. For a recorded iteration, we calculate the distances *D*_*i*_0_,*i*_ between the viewpoint *i*_0_ and all other bins *i* ≠ *i*_0_, and sample the duration time 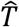 of the current state by exponential distribution. If 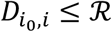, we increase *T_i_* by 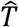. Finally, *M_i,j_, L_i_, R_i_* are updated until the next recorded iteration comes (updated 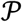 times). The above process is performed iteratively and stops after enough iterations. The final *T_i_* for *i* ∈ [1, *I*] is normalized by 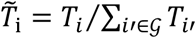, where 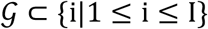 is the set of bins involved in normalization, *e.g*. bins containing genes. Simulated figures are generated by drawing bar of height 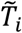 at each bin *i*.

### Parameter optimization

We used permeability *p* = 0.1125 and *q* = 1, separation *S* = 750 kb, and processivity *λ* = 2.5 Mb. Simulation is initialized at the state without any loaded Cohesin, i.e. *M_i,j_* = 0 for 1 ≤ *i* ≤ *j* ≤ *I* and *L_i_* = *R_i_* = 0 for 1 ≤ *i* ≤ *I*. We performed each simulation with *L* = 1.5 × 10^9^ iterations, skipped the first 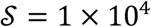 iterations, and then sampled every 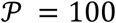 iterations. Distance between bins connected by Cohesin is set by *C* = 0. Bins with distance no more than *ℛ* = 0 to the viewpoint are considered to have formed long-distance chromatin interactions.

### Genome-wide insulator analyses

#### Define promoter and its activities

To investigate whether CBS located between enhancer and promoter functions as an insulator in the genome-wide scale, we define transcription start sites (TSS) as promoters for each gene. Specifically, for genes with multiple TSSs, we consider that they have multiple promoters. Promoter activities were assayed by RNA-seq experiments with expression levels as FPKM values. In cases of different transcripts shared the same TSS, the expression levels are defined as the sum of FPKM from all of the transcripts with the same TSS.

#### Define insulator and its strength

We use FIMO (version 5.0.1) to call putative CBSs in both orientations of the hg19 human genome assembly by default parameters with CTCF binding position weight matrices (PWM), which define the probability *P_n,i_* for the observed nucleotides *n* ∈ {*A, C, G, T*} at position *i* ∈ [1, 42] of the CTCF motif [12, 52].

We performed ChIP-seq experiments with a specific antibody against CTCF for each of the three cell lines as described above. In addition, we also downloaded CTCF ChIP-seq data from the ENCODE datasets with accession numbers: GSM923425, GSM1003581, and GSM1003578 for A549 cells; GSM958728, GSM935611, and GSM733771 for GM12878 cells; GSM958749, GSM1022652, and GSM945853 for HCT 116 cells; GSM958735, GSM822285, and GSM733684 for HeLa cells; wgEncodeEH000135, GSM749683, and GSM733743 for HepG2 cells; GSM958736, GSM733636, GSM733674 for NHEK cells [40]. CTCF peaks are called by MACS (version 1.4.2) with a cutoff *P* value of 10^−5^, and enrichment is assigned to each peak automatically.

We extend each CTCF peak by 50 bp in both directions. For putative CBS intersecting with at least one extended CTCF peak, we assign it to the closest CTCF peak. These may result in that several CBSs are assigned to the same CTCF peak. In these cases, enrichment of CTCF proteins with a ChIP-seq peak is equally divided to all CBSs associated with it. For CBS not intersecting with any extended CTCF peak, it is abandoned.

We assume that each CTCF peak should contain at least one CBS and develop the following method to find the most likely CBS within the CTCF peak that does not intersect with any putative CBS. Thus, for CTCF peak (*d_l_, d_r_*), whose extension (*d_l_* − 50, *d_r_* + 50) does not intersect with any CBS, we define the forward 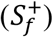 and reverse 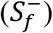 scores of the segment (*f, f* + 41), *f* ∈ [*d_l_* − 50 − 42 + 1, *d_r_* + 50] as

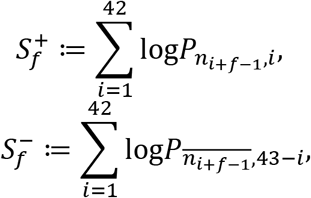

where *n*_*i*+*f*−1_ is the nucleotide at position *i* + *f* − 1 of hg19, and 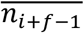 is the complement (A to T, T to A, G to C, and C to G) of *n*_*i*+*f*−1_. Let

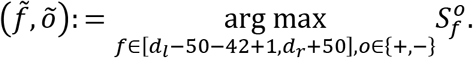

Then 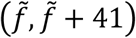 is considered as CBS in orientation *õ*, though it is not called by FIMO, because among all segments intersecting with the extended CTCF peak (*d_l_* − 50, *d_r_* + 50), it fits the CBS position weighted matrix best.

Finally, insulators are defined as CBSs, and their insulation efficiencies are the assigned CTCF protein enrichments.

#### Define promoter-enhancer pairs

We perform ChIP-seq experiments with a specific antibody against H3K27ac and define active enhancers as H3K27ac peaks (called by MACS version 1.4.2). We pair each promoter (TSS) with at least one enhancer, and vice versa. Thus, there is no unpaired promoter or enhancer. We then use a pairing method that minimizes the total distance of all pairs. The position of TSS is indicated by a single coordinate *t* on single chromosomes of the hg19 assembly. The enhancer peak range is indicated by an interval (*e^l^, e^r^*) from the left edge to the right edge of the H3K27ac ChIP-seq peak on the same chromosome as the TSS. Distances between TSS *t* and enhancer peak (*e^l^, e^r^*) is defined as |*t* − *e^m^*|, with *e^m^* = (*e^l^* + *e^r^*)/2 as the middle point of the peak.

For TSSs of two promoters where *t* < *t′* and two enhancers where *e^m^* < *e^m′^*, there is always |*t* − *e^m^*| + |*t′* − *e^m′^*| ≤ |*t′* − *e^m′^*| + |*t′*, − *e^m^*|. In fact, let 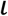 be any position coordinate between max(*t, e^m^*) and min(*t′, e^m′^*). Then 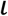 separates not only *t* and *e^m′^*, but also *e^m^*and *t′*. As a result, 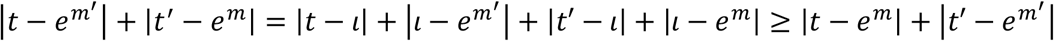. Thus, we do not consider crossing enhancer-promoter pairs like (*t, e^m′^*) and (*t′, e^m^*).

The pairing method with the minimal total distance of all pairs can be developed by dynamic programming. Suppose there are TSSs *t*_1_ < *t*_2_ < ⋯ < *t_U_* and enhancers 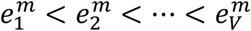. For *i* ∈ [1, *U*] and *j* ∈ [1, *V*], let *E_i,j_* be the minimal total distance among all non-crossing pairing methods between *t*_1_, *t*_2_, −, *t_i_* and 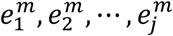. Obviously,

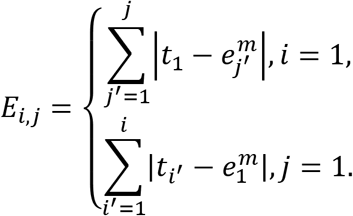

For *i,j* > 1, *t_i_* must pair with 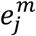 in any non-crossing pairing method between *t*_1_, *t*_2_, ⋯, *t_i_* and 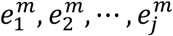. Otherwise, they respectively pair with 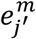 and *t_i′_* for certain 1 ≤ *j′* < *j* and 1 ≤ *i′* < *i*, which conflicts with the non-crossing assumption. Now we have three cases as following:

1. If *t_i_* pairs with 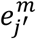 for certain 1 ≤ *j′* < *j*, then 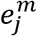 only pairs with *t_i_*, because otherwise, the pairing method is crossing. Thus, removing the pair 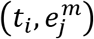 will lead to a non-crossing pairing method between *t*_1_, *t*_2_, ⋯, *t_i_* and 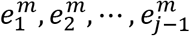.
2. If *t_i_* does not pair with 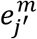 for any 1 ≤ *j′* < *j*, and 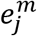 pairs with for certain 1 ≤ *i′* < *i*, then removing the pair 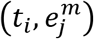 will lead to a non-crossing pairing method between *t*_1_, *t*_2_, ⋯, *t*_*i*−1_ and 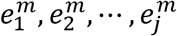.
3. If *t_i_* does not pair with 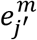 for any 1 ≤ *j′* < *j*, and 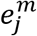 does not pair with for any 1 ≤ *i′* < *i*, then removing the pair 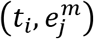 will lead to a non-crossing pairing method between *t*_1_, *t*_2_, ⋯, *t*_*i*−1_ and 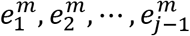.

Therefore, for *i,j* > 1,

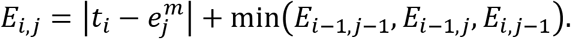

Thus, one may recursively calculate *E_i,j_* ∀*i* ∈ [1, *U*], *j* ∈ [1, *V*] by the above two equations during which we record the minimum among *E*_*i*−1,*j*−1_, *E*_*i*−1,j_, *E*_*i,j*−1_ in order to backtrack all promoter-enhancer pairs. Specifically, we start from *E_U,V_*. Suppose that we are now at *E_i,j_* for *i, j* > 1. First, we pair *t_i_* with 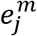, and move to the minimum of *E*_*i*−1,*j*−1_, *E*_*i*−1,j_, *E*_*i,j*−1_. If the minimum is not unique, we choose *E*_*i*−1,*j*_, then *E*_*i*−1,*j*−1_, and finally *E*_*i,j*−1_. This process is performed iteratively until we arrive at *E_i,j_* with *i* = 1 or *j* = 1. Assuming without loss of generality that *i* = 1, we then pair all 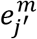 for *j′* ∈ [1, *j*] with *t*_1_.

#### Filter promoter-enhancer pairs

Since H3K27ac may also mark some promoters, if promoters and enhancers are too close, enhancers may actually be promoters. Thus, we filter all promoter-enhancer pairs (*t, e^m^*) with |*t* − *e^m^*| < 1 kb.

#### Define insulator score

Assuming without loss of generality that *t* < *e^m^* in the promoter-enhancer pair (*t, e^m^*), we say that a CBS (*f, f* + 41) insulates the pair (*t, e^m^*) if and only if *t* + 400 bp < *f^m^* < *e^m^* − 400 bp, where *f^m^* = *f* + 41/2 is the middle of CBS. Let Ξ be the enrichment of H3K27ac in the enhancer *e^m^*, and Φ be the enrichment of CTCF in the CBS element insulating the pair (*t, e^m^*). Then the activation power of enhancer *e^m^* for promoter *t* is defined as Ξ/(Φ + 1). We add one on the denominator in order to handle vanishing Φ. For those promoters paired with several enhancers, the total activation power for promoters is the sum of the activation power of all enhancers pairing it. Finally, we define the insulator score as the reverse of the activation power.

#### Visualization

Denote the sorted promoters by *t*_1_, *t*_2_, ⋯, *t_U_*. Let *δ_i_* be the insulator score for promoter *t_i_*. The red line in each panel is generated through fitting *δ_i_* and *i* by linear least squares regression. Define 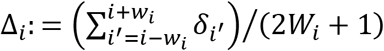, where *W_i_* = min(100, *i, U* − *i*), *i.e*. Δ_*i*_ is the average of *δ_i_* in a sliding window of length 2*W_i_* + 1. The fluctuating line is a plot between *i* and *Δ_i_*. The limitations of the vertical axis are normalized to [0,1]. Because FPKM monotonically increases with *i*, the vertical axis actually indicates the activities of promoters.

## ACKNOWLEDGMENTS

This work was supported by grants from NSFC (31630039, 91640118, and 31470820) and MOST (2017YFA0504203, 2018YFC1004504) to Q.W. Q.W. is a Shanghai Subject Chief Scientist.

---

**Figure S1.**
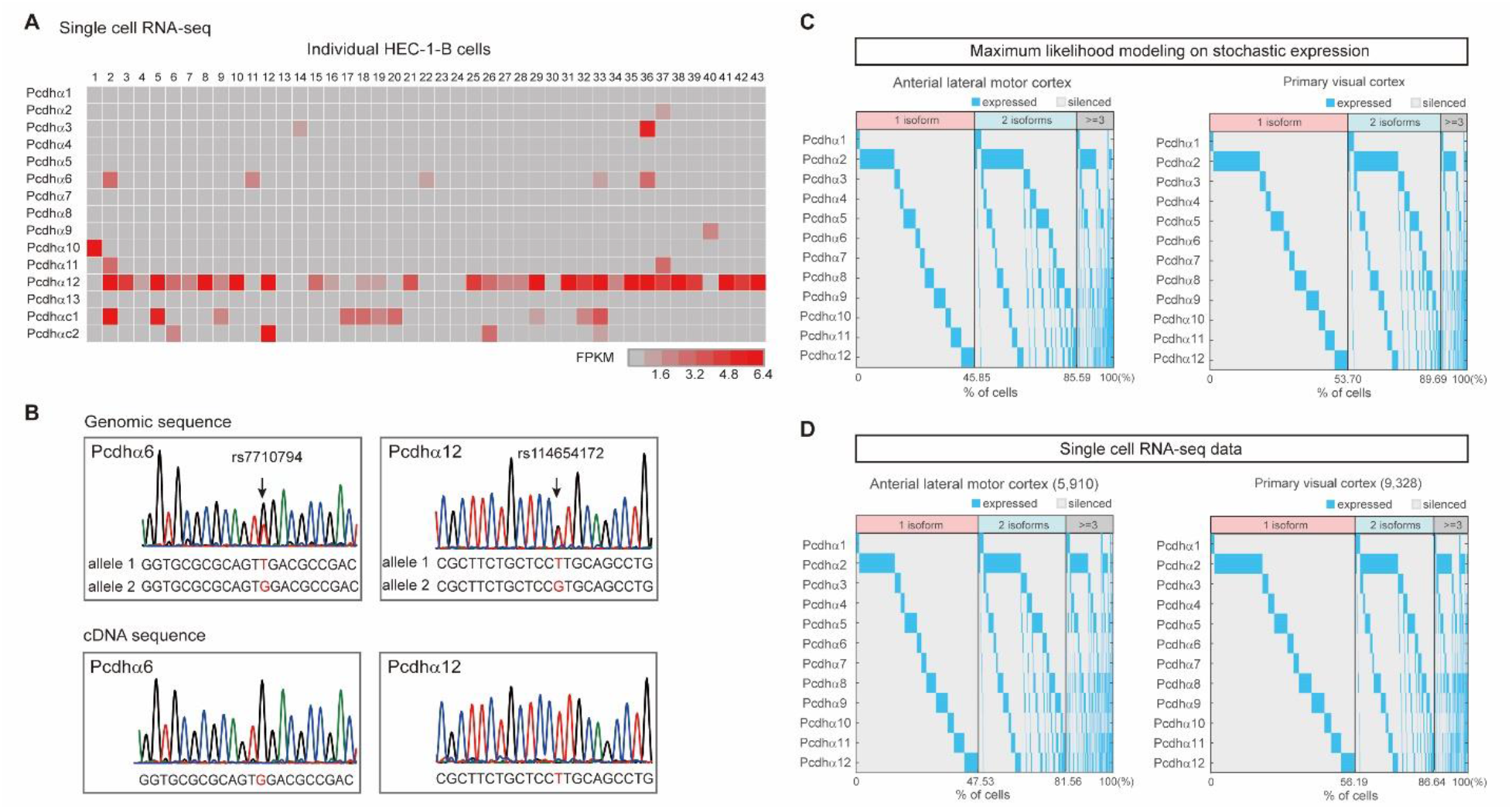
Stochastic and monoallelic expression of *Pcdhα* genes in single cells. (**A**) Shown are stochastic expression patterns of the *Pcdhα* genes in individual HEC-1-B cells by single-cell RNA-seq. (**B**) Shown are Sanger sequencing traces for genomic DNA and cDNA. Note that two SNPs are detected in *α6* and *α12* variable exons, but only one allele is expressed in single HEC-1-B cells. (**C**) Maximum likelihood prediction of *Pcdhα* gene expression patterns in single cells based on stochastic expression assumption. (**D**) Stochastic expression patterns of the *Pcdhα* genes in single cells from different mouse cortical regions, with most cells express one or two isoforms. Note that the expression patterns are similar between prediction and experimental data.

**Figure S2.**
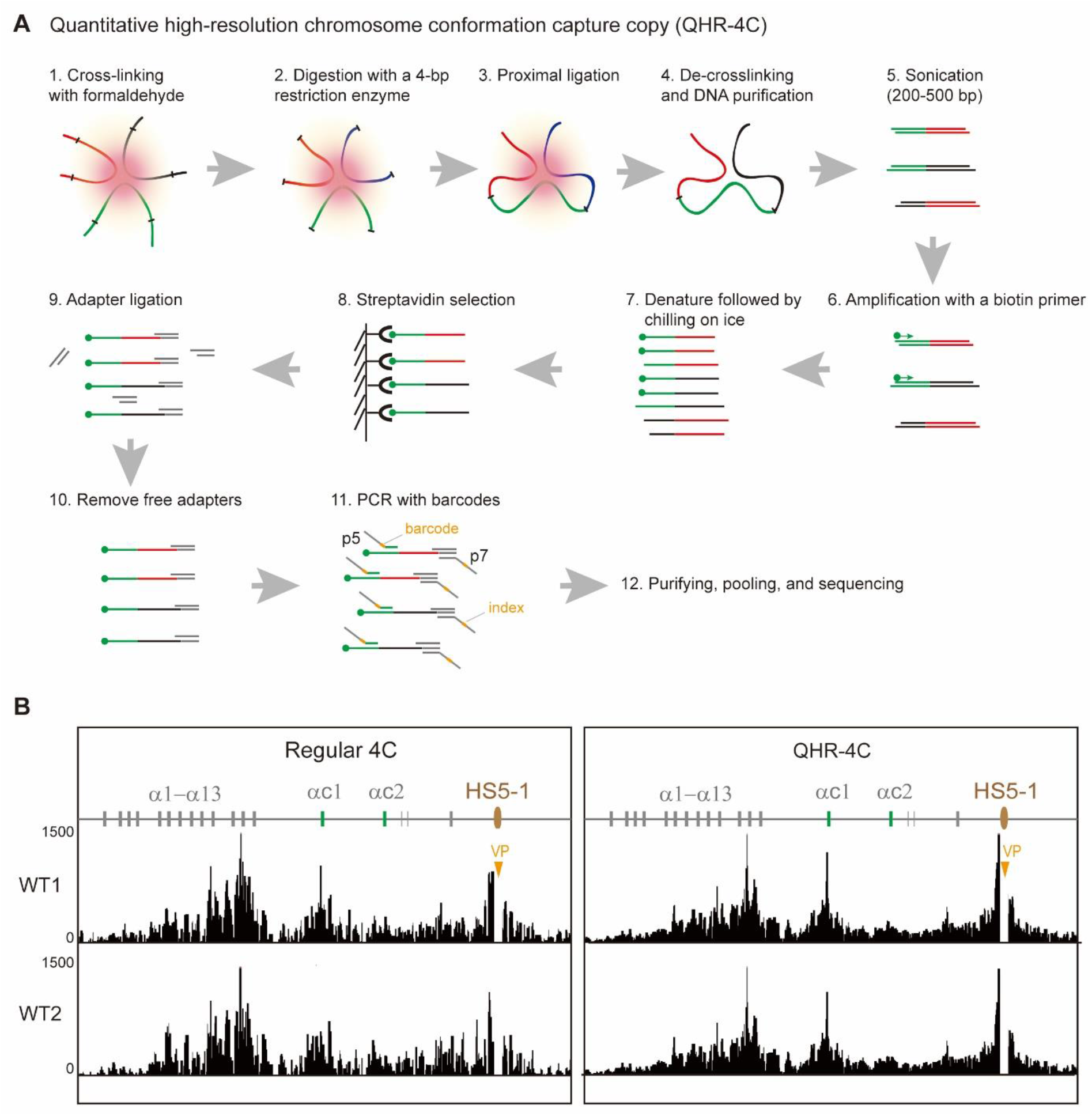
A sensitive QHR-4C method for capturing chromosome conformation. (**A**) Schematic of the QHR-4C method for detecting long-range chromatin interaction profiles of a viewpoint (see QHR-4C section in Materials and Methods above for details). (**B**) Shown are a comparison between the regular- and QHR-4C methods of chromatin interaction profiles. VP: viewpoint.

**Figure S3.**
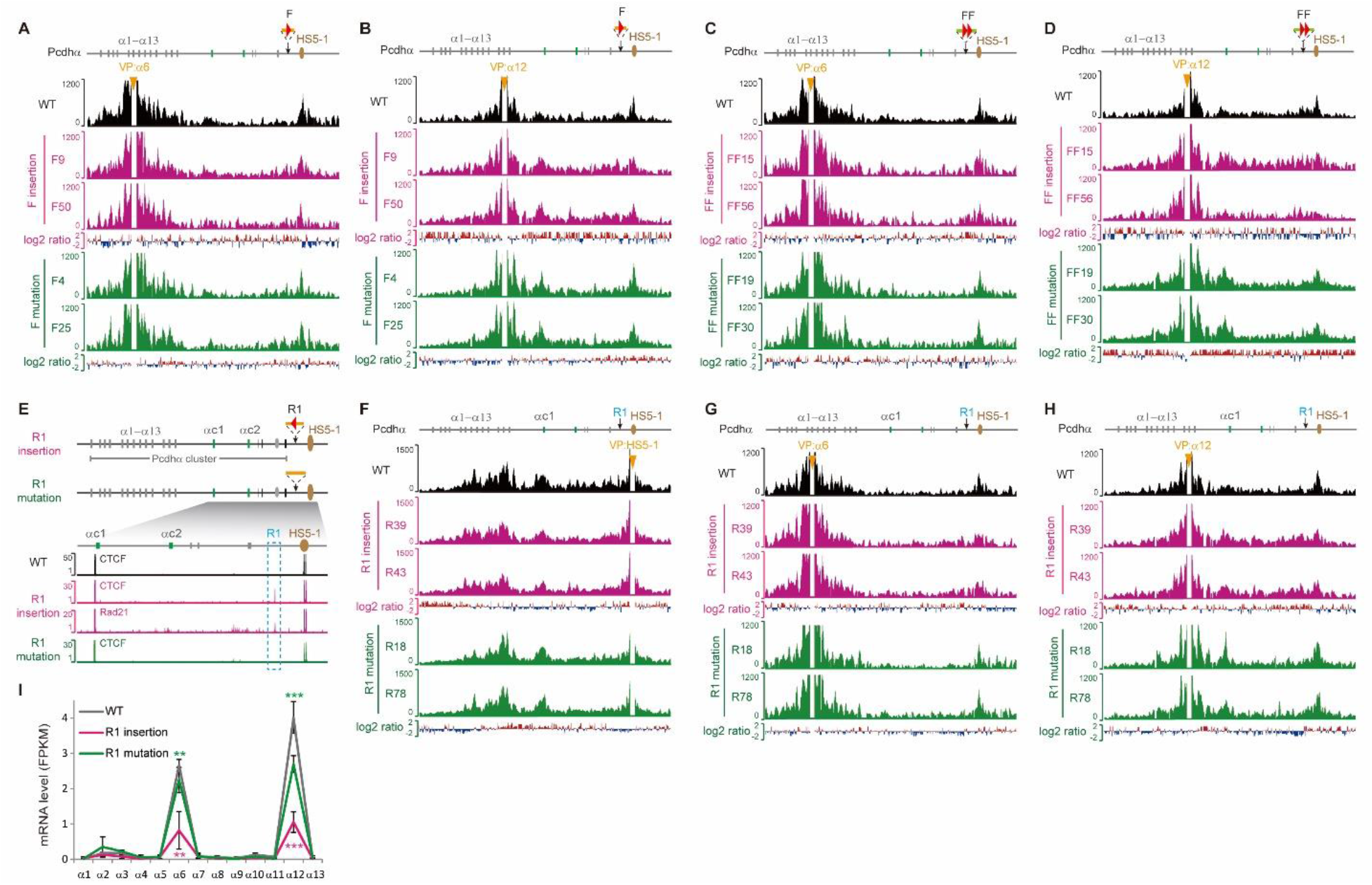
Both forward and reverse CBSs inserted between the *Pcdhα* cluster and its downstream *HS5-1* enhancer function as insulators. (**A** and **B**) 4C interaction profiles with *a6* or *a12* as a viewpoint in individual CRISPR single-cell clones with insertions of one forward CBS or its mutation. (**C** and **D**) 4C interaction profiles with *a6* or *a12* as a viewpoint in individual CRISPR single-cell clones with insertions of two forward CBSs or their mutations. (**E**) Schematic showing the insertion of the reverse CBS (“R1”) or its mutation into the location between *Pcdhα* cluster and its downstream *HS5-1* enhancer. CTCF and Rad21 ChIP-seq confirmed their binding to the inserted CBS. For clarity, only regions around insertions are shown. (**F**) 4C profiles with *HS5-1* as a viewpoint for CRISPR single-cell clones with the insertion of wildtype CBS or its mutation. (**G**) 4C profiles with the *a6* promoter as a viewpoint. (**H**) 4C profiles with the *a12* promoter as a viewpoint. (**I**) Gene expression patterns in CRISPR single-cell clones as measured by RNA-seq. Data as mean ± SD, \*\**p* < 0.01, \*\*\**p* < 0.001; one-tailed Student’s *t* test.

**Figure S4.**
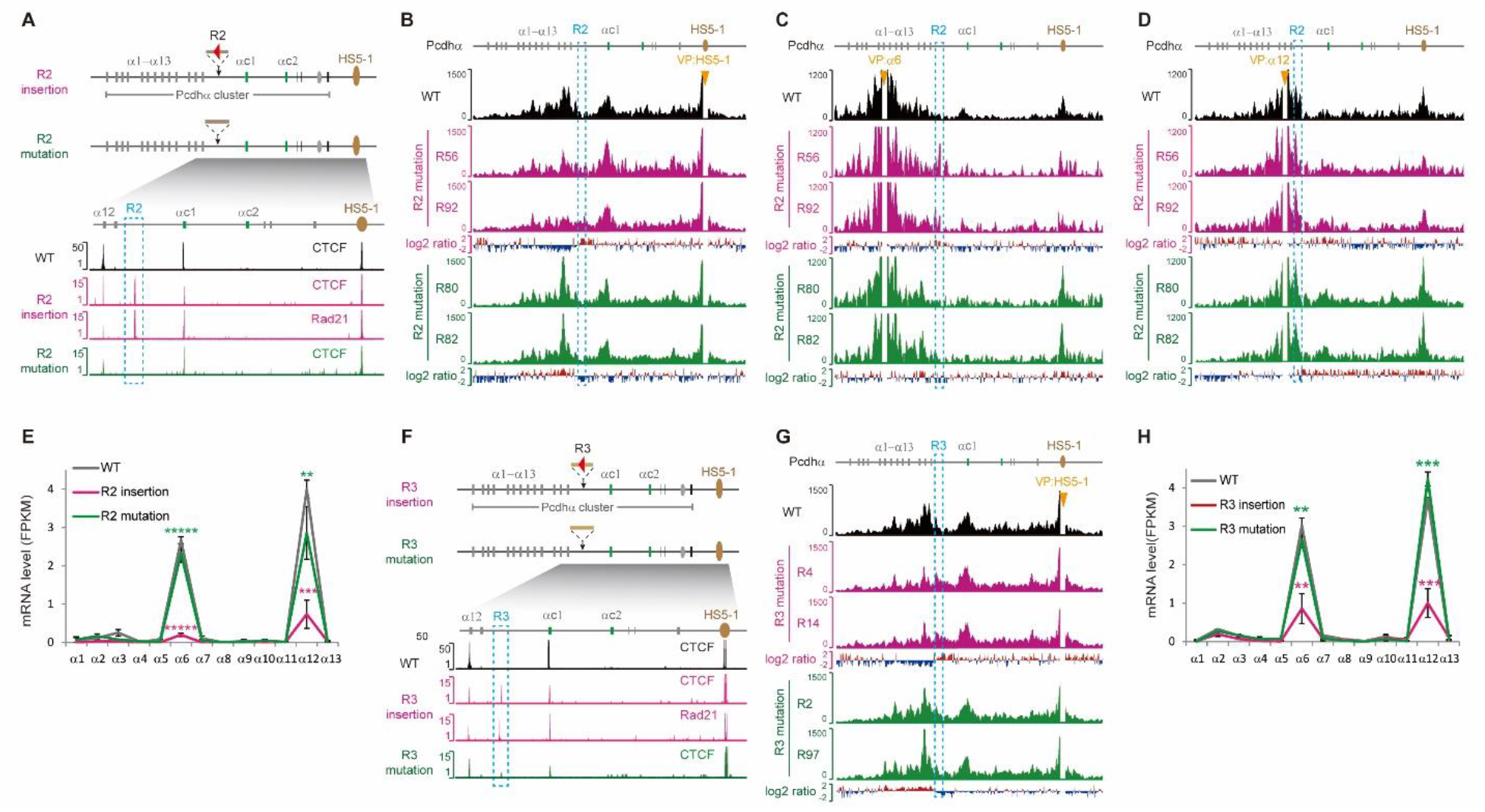
Reverse CBSs inserted between *Pcdh α13* and *αc1* function as an insulator for its upstream genes. (**A**) Schematic showing the insertion of the reverse CBS (“R2”) into the location between *α13* and *αc1* within the *Pcdhα* cluster. CTCF and Rad21 ChIP-seq are shown below. (**B**) 4C profiles with *HS5-1* as a viewpoint. (**C**) 4C profiles with the *α6* promoter as a viewpoint. (**D**) 4C profiles with the *α12* promoter as a viewpoint. (**E**) Gene expression levels in CRISPR single-cell clones as measured by RNA-seq. (**F**) Schematic showing the insertion of the reverse CBS (“R3”) into the location between *α13* and *αc1* within the *Pcdhα* cluster. CTCF and Rad21 ChIP-seq are shown below. (**G**) 4C profiles with *HS5-1* as a viewpoint. (**H**) Gene expression levels measured by RNA-seq. Data as mean ± SD, \*\**p* < 0.01, \*\*\**p* < 0.001, \*\*\*\*\**p* < 0.00001; one-tailed Student’s *t* test.

**Figure S5.**
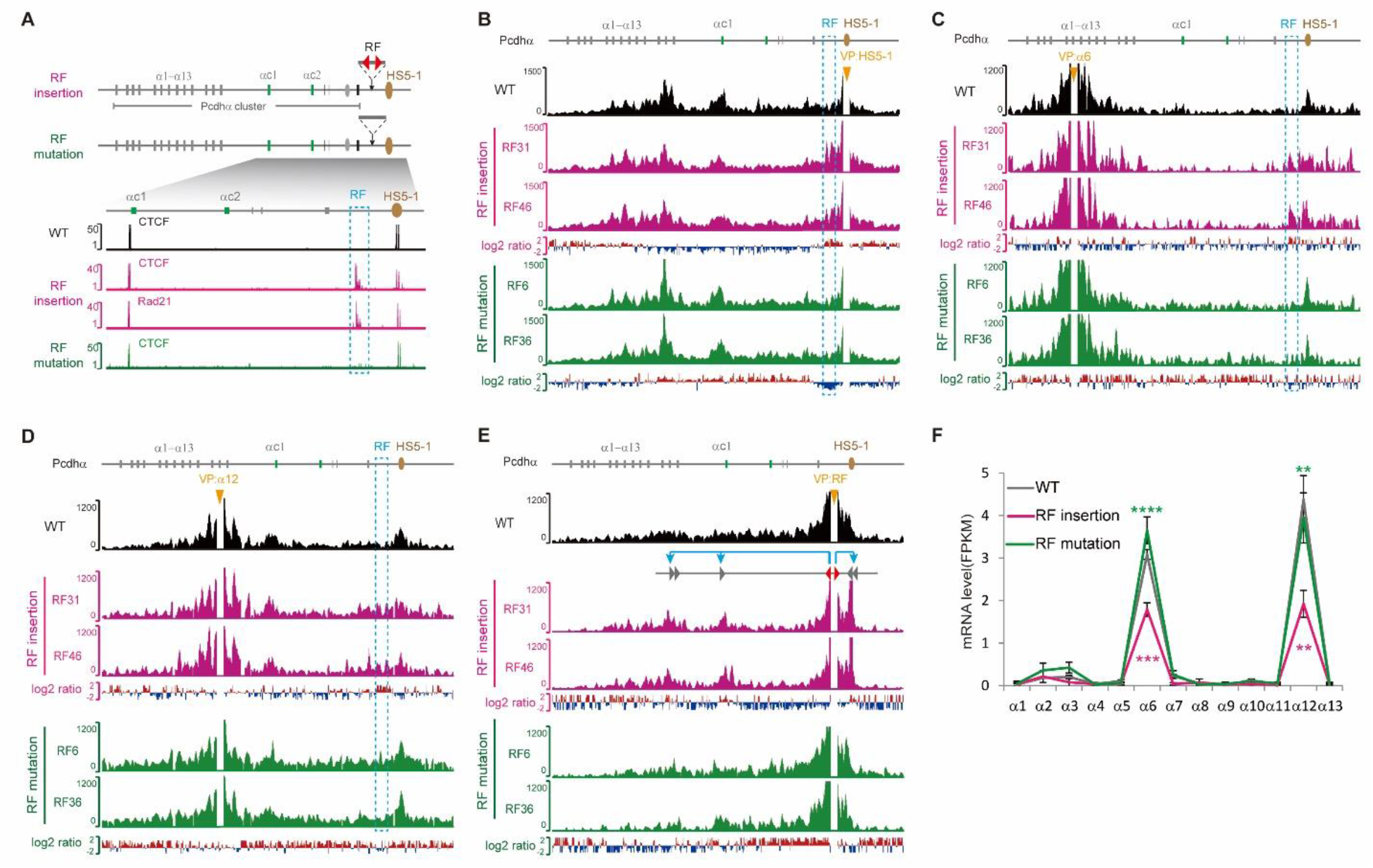
Reverse-forward CBS pair as an insulator for the *Pcdhα* genes. (**A**) Schematic showing the insertion of reverse-forward CBSs (“RF”) into the location between the *Pcdhα* cluster and its downstream *HS5-1* enhancer. CTCF and Rad21 ChIP-seq are shown below. (**B**) 4C profiles with *HS5-1* as a viewpoint. (**C**) 4C profiles with the *a6* promoter as a viewpoint. (**D**) 4C profiles with the *a12* promoter as a viewpoint. (**E**) 4C profiles with the inserted region as a viewpoint. (**F**) Gene expression levels measured by RNA-seq. Data as mean ± SD, \*\**p* < 0.01, \*\*\**p* < 0.001, \*\*\*\**p* < 0.0001; one-tailed Student’s *t* test.

**Figure S6.**
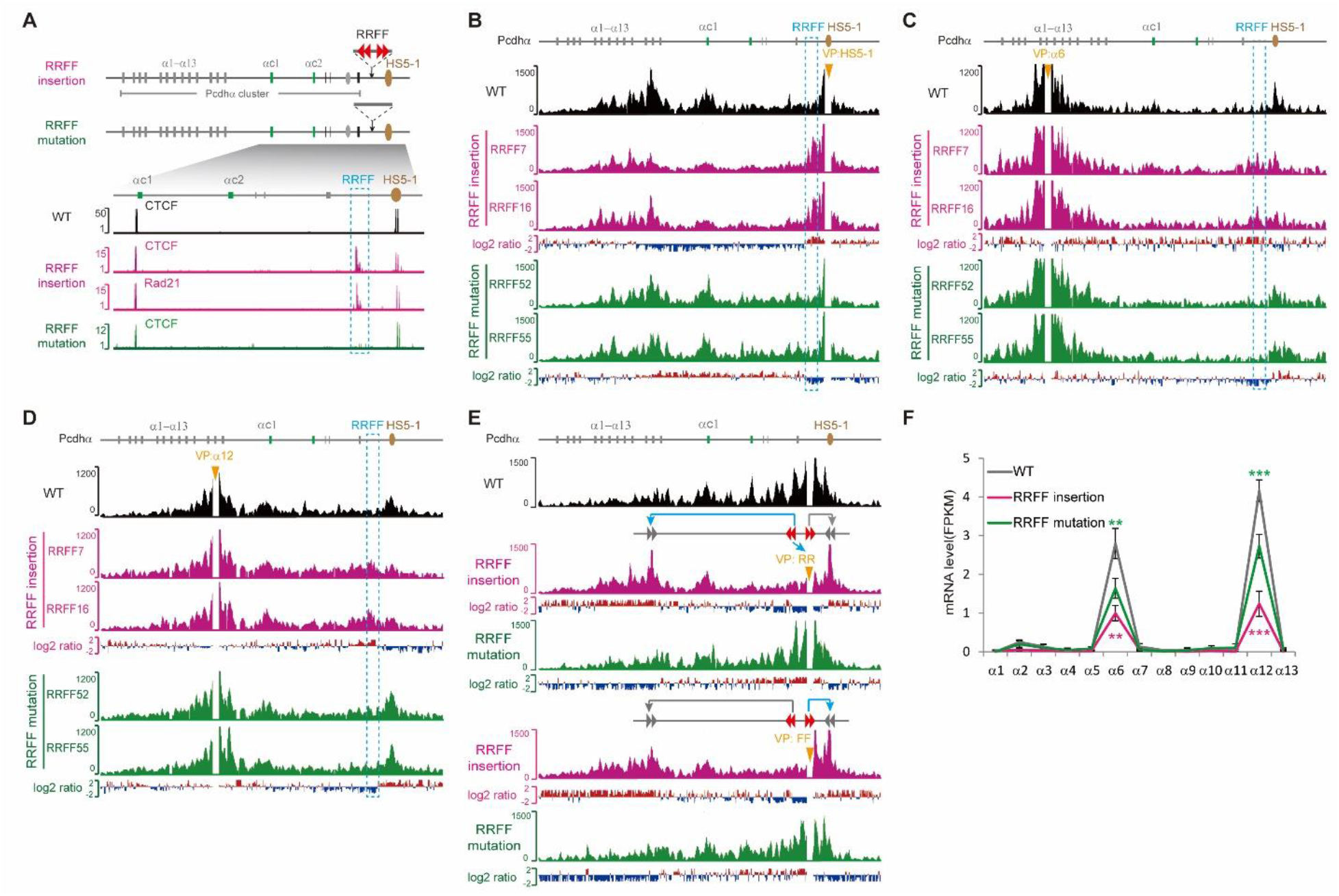
Reverse-forward tandem CBS pairs as an insulator for the *Pcdhα* genes. (**A**) Shown are the insertions of reverse-forward CBSs (“RRFF”) into the location between the *Pcdhα* cluster and its downstream *HS5-1* enhancer. CTCF and Rad21 ChIP-seq are shown below. (**B**) 4C profiles with *HS5-1* as a viewpoint. (**C**) 4C profiles with the *a6* promoter as a viewpoint. (**D**) 4C profiles with the *a12* promoter as a viewpoint. (**E**) 4C profiles with the inserted region as a viewpoint. (**F**) Gene expression levels measured by RNA-seq. Data as mean ± SD, \*\**p* < 0.01, \*\*\**p* < 0.001; one-tailed Student’s *t* test.

**Figure S7.**
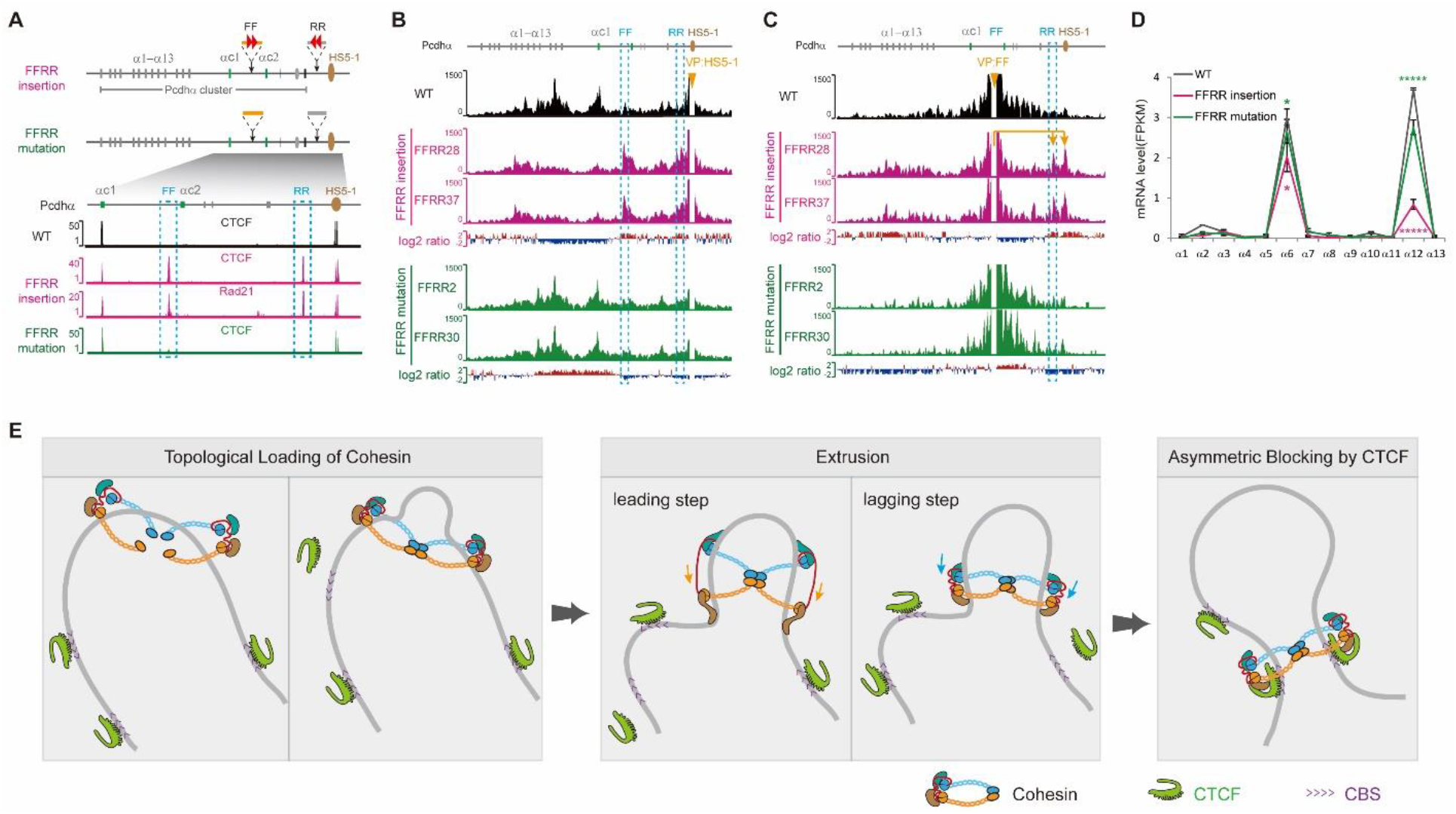
Forward-reverse CBS pairs as an insulator for the upstream *Pcdhα* genes. (**A**) Schematic showing the insertions of forward (“FF”) and reverse (“RR”) CBSs into the *Pcdhα* cluster. CTCF and Rad21 ChIP-seq are shown below. (**B**) 4C profiles with *HS5-1* as a viewpoint. (**C**) 4C profiles with the inserted forward CBSs as a viewpoint in the CRISPR single-cell clones. (**D**) Gene expression levels measured by RNA-seq. Data as mean ± SD, \**p* < 0.05, \*\*\*\*\**p* < 0.00001; one-tailed Student’s *t* test. (**E**) A “double clamp” extrusion model for Cohesin-mediated loop formation. We proposed that, upon topologically loading, each ring of the Cohesin dimer embracing a dsDNA within the lumen. The two Cohesin rings interact with each other and slide along dsDNA in opposite directions. The extrusion process can be divided into a leading step and a lagging step, which depends on ATP hydrolyses. Extrusion is continuous until blocked by oriented CBSs bound with CTCF which anchors Cohesin in an orientation-dependent manner.

**Figure S8.**
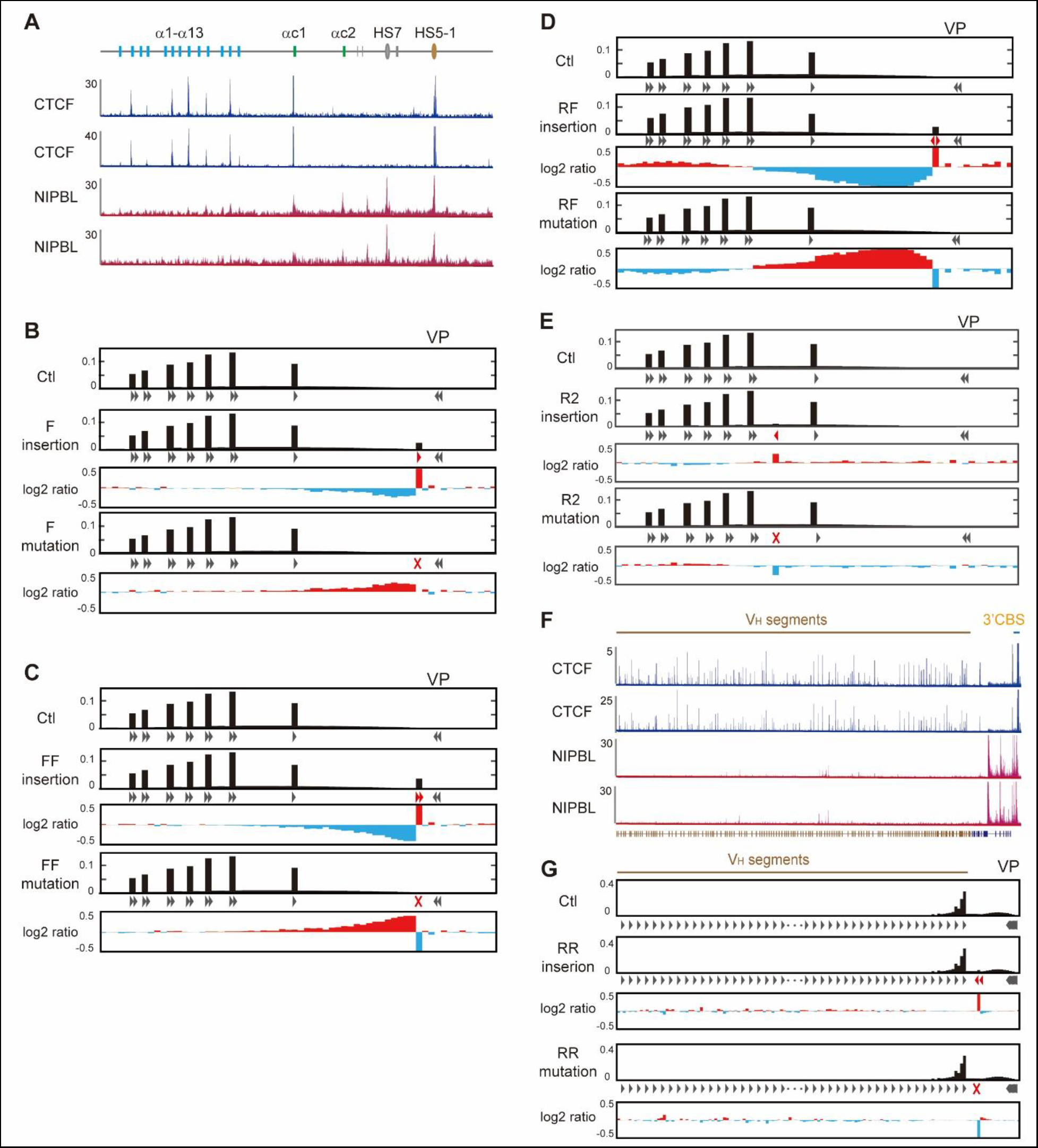
Simulations of the chromatin looping interaction profiles upon CBS insertions or their mutations in the *Pcdh* or *Igh* clusters. (**A**) ChIP-seq tracks of CTCF and NIPBL at the *Pcdhα* cluster in HEC-1-B cells. (**B-D**) Simulation of interaction profiles with *HS5-1* as a viewpoint for CRISPR single-cell clones with the insertion of one forward CBS (**B**) or two forward CBSs (**C**) or a pair of divergent CBSs (**D**) or a reverse CBS in a different location (**E**) and their corresponding mutations, respectively. (**F**) ChIP-seq tracks of CTCF and NIPBL at the *Igh* cluster in pro-B cells. (**G**) Simulation of interaction profiles with *3’CBS* as a viewpoint in pro-B cells.

**Figure S9.**
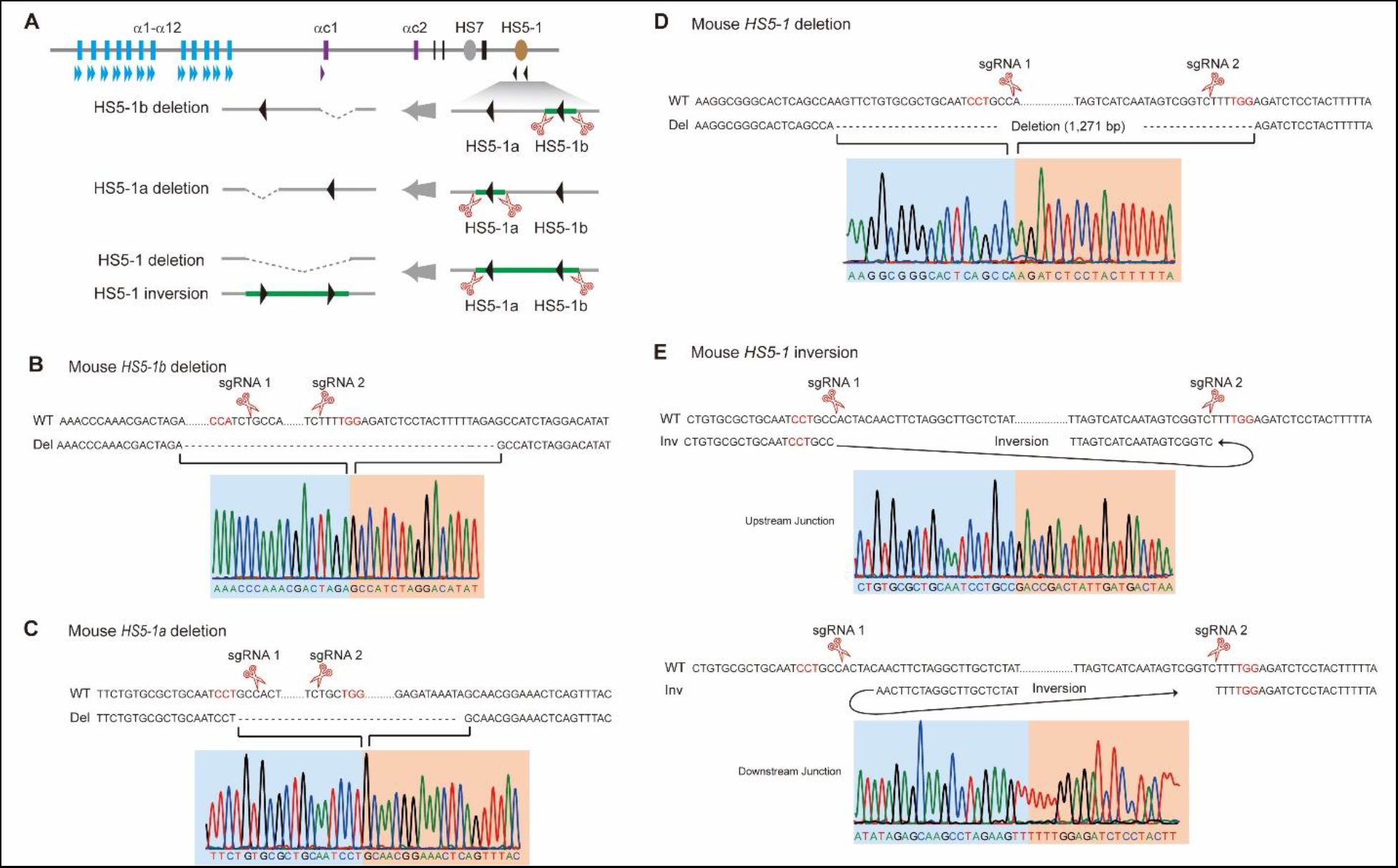
Sanger sequencing traces for the deletion or inversion mice generated by DNA-fragment editing. (**A**) Schematic of the deletion and inversion mouse lines. (**B-D**) Shown are Sanger sequencing traces of deletion junctions for the *HS5-1b* (**B**) or *HS5-1a* (**C**) or *HS5-1* (**D**) homozygous mice generated by DNA-fragment editing. (**E**) Sanger sequencing traces of genotyping for the upstream and downstream junctions of *HS5-1* homozygous inversion mice. The PAM sites are highlighted.

**Figure S10.**
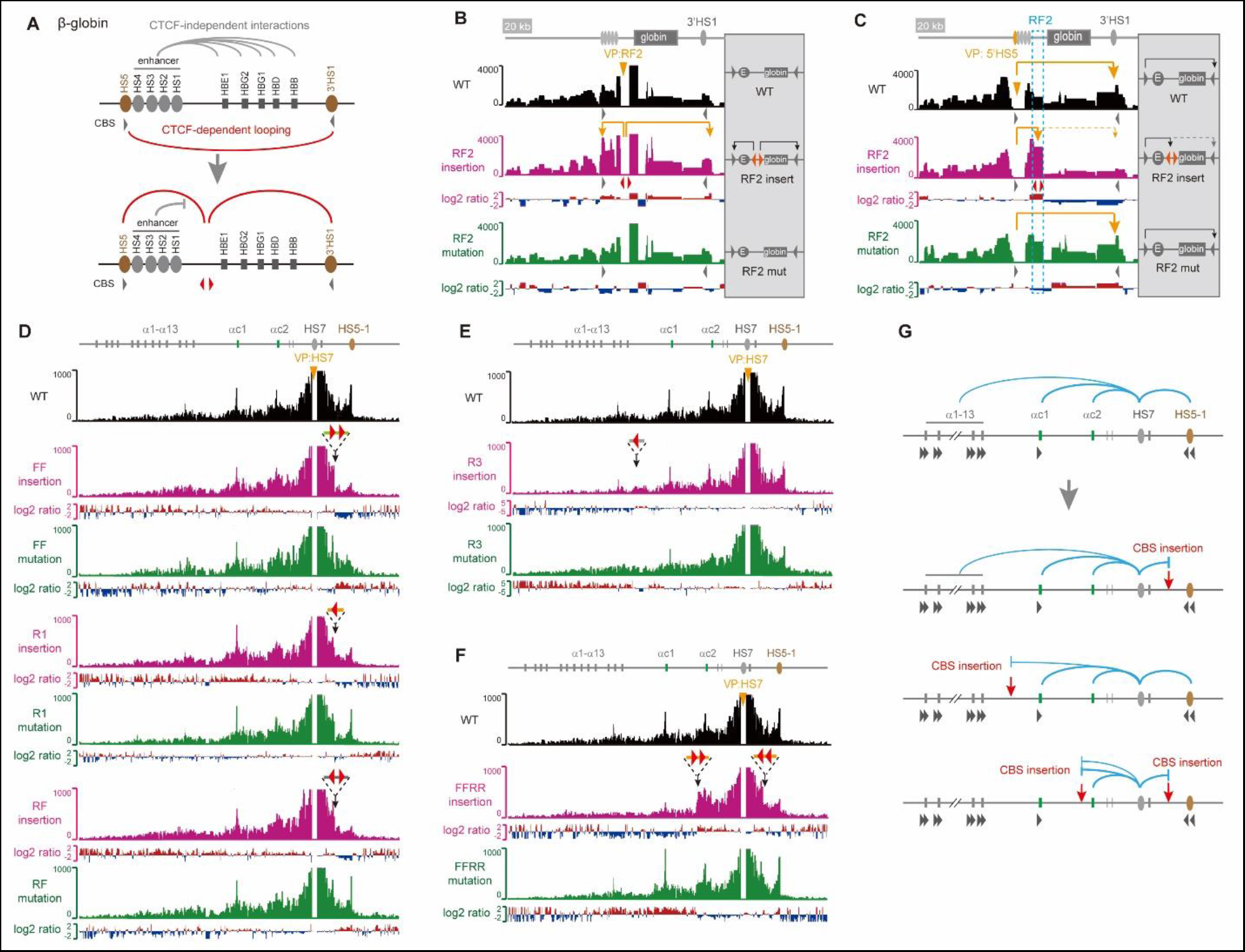
Long-distance chromatin interaction profiles for enhancers with no CBS. (**A**) Schematic of the chromatin interaction profiles upon insertion of a divergent CBS pair in the *β-globin* cluster. (**B**) 4C profiles with the inserted CBS as a viewpoint in the *β-globin* cluster. (**C**) 4C profiles with 5’HS5 CBS as a viewpoint in the *β-globin* cluster. (**D**) 4C profiles with *HS7* as a viewpoint revealed a significant decrease of chromatin looping interactions with *HS5-1* upon the insertion of tandem forward CBSs or single reverse CBS or a pair of reverse-forward CBSs. Note a sharp decrease of interactions as shown by log2 ratio at the exact insertion site (probably because the bound CTCF blocks Cohesin sliding at the site) and the rescue upon the CBS mutations. (**E**) 4C profile with *HS7* as a viewpoint revealed a significant decrease of chromatin looping interactions with the alternate *Pcdhα* genes upon the insertion of a reverse CBS into the location between *Pcdh a13* and *ac1*. Note that CBS mutation rescues the blocking effects. (**F**) 4C profile with *HS7* as a viewpoint revealed a significant decrease of chromatin looping interactions beyond both upstream and downstream insertion sites. Note that CBS mutations rescue the blocking effects. (**G**) Schematic of the altered chromatin interaction profiles upon various insertions.

**Figure S11.**
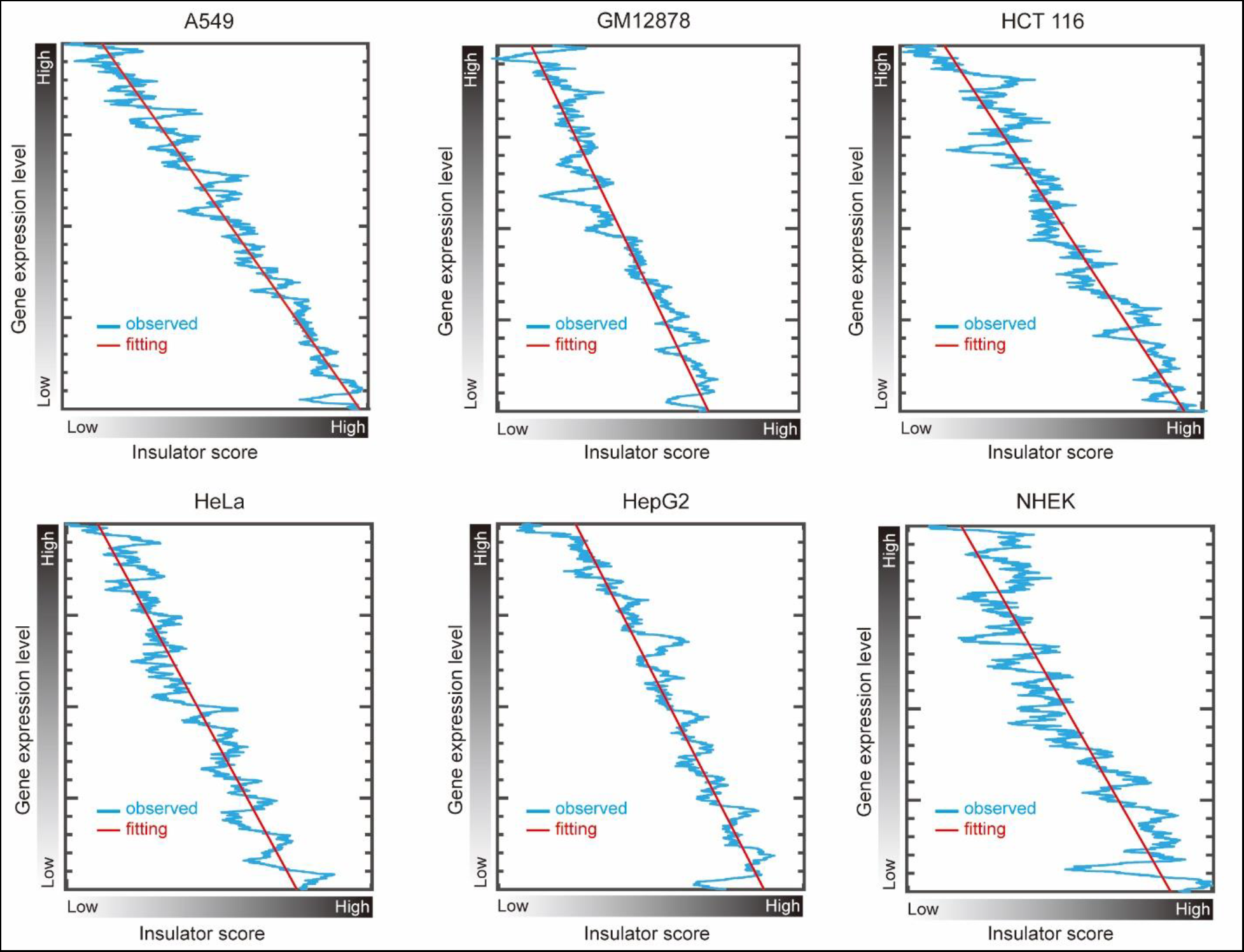
Negative correlations of gene expression levels with insulator scores in 6 additional cell lines. The insulator score is defined by CTCF dynamics and enhancer strength as described in Materials and Methods.

## REFERENCES

1. Muller, H.J., Types of visible variations induced by X-rays in Drosophila. Journal of Genetics, 1930. 22(3): p. 299–334.

2. Phillips-Cremins, J.E. and V.G. Corces, Chromatin insulators: linking genome organization to cellular function. Mol Cell, 2013. 50(4): p. 461–474.

3. Dekker, J. and L. Mirny, The 3D genome as moderator of chromosomal communication. Cell, 2016. 164(6): p. 1110–1121.

4. Furlong, E.E.M. and M. Levine, Developmental enhancers and chromosome topology. Science, 2018. 361(6409): p. 1341–1345.

5. Grosveld, F., G.B. van Assendelft, D.R. Greaves, and G. Kollias, Position-independent, high-level expression of the human beta-globin gene in transgenic mice. Cell, 1987. 51(6): p. 975–985.

6. Chung, J.H., M. Whiteley, and G. Felsenfeld, A 5’ element of the chicken beta-globin domain serves as an insulator in human erythroid cells and protects against position effect in Drosophila. Cell, 1993. 74(3): p. 505–514.

7. Bell, A.C., A.G. West, and G. Felsenfeld, The protein CTCF is required for the enhancer blocking activity of vertebrate insulators. Cell, 1999. 98(3): p. 387–396.

8. Ghirlando, R. and G. Felsenfeld, CTCF: making the right connections. Genes Dev, 2016. 30(8): p. 881–891.

9. Hansen, A.S., I. Pustova, C. Cattoglio, R. Tjian, and X. Darzacq, CTCF and cohesin regulate chromatin loop stability with distinct dynamics. Elife, 2017. 6.

10. Xu, D., R. Ma, J. Zhang, Z. Liu, B. Wu, J. Peng, Y. Zhai, Q. Gong, Y. Shi, J. Wu, Q. Wu, Z. Zhang, and K. Ruan, Dynamic nature of CTCF tandem 11 zinc fingers in multivalent recognition of DNA as revealed by NMR spectroscopy. J. Phys. Chem. Lett., 2018. 9(14): p. 4020–4028.

11. Rao, S.S.P., M.H. Huntley, N.C. Durand, E.K. Stamenova, I.D. Bochkov, J.T. Robinson, A.L. Sanborn, I. Machol, A.D. Omer, E.S. Lander, and E.L. Aiden, A 3D map of the human genome at kilobase resolution reveals principles of chromatin looping. Cell, 2014. 159(7): p. 1665–1680.

12. Guo, Y., Q. Xu, D. Canzio, J. Shou, J. Li, D.U. Gorkin, I. Jung, H. Wu, Y. Zhai, Y. Tang, Y. Lu, Y. Wu, Z. Jia, W. Li, M.Q. Zhang, B. Ren, A.R. Krainer, T. Maniatis, and Q. Wu, CRISPR inversion of CTCF sites alters genome topology and enhancer/promoter function. Cell, 2015. 162(4): p. 900–910.

13. Rudan, M.V., C. Barrington, S. Henderson, C. Ernst, D.T. Odom, A. Tanay, and S. Hadjur, Comparative Hi-C reveals that CTCF underlies evolution of chromosomal domain architecture. Cell Rep, 2015. 10(8): p. 1297–1309.

14. de Wit, E., E.S.M. Vos, S.J.B. Holwerda, C. Valdes-Quezada, M.J.A.M. Verstegen, H. Teunissen, E. Splinter, P.J. Wijchers, P.H.L. Krijger, and W. de Laat, CTCF binding polarity determines chromatin looping. Mol Cell, 2015. 60(4): p. 676–684.

15. Hnisz, D., A.S. Weintraub, D.S. Day, A.L. Valton, R.O. Bak, C.H. Li, J. Goldmann, B.R. Lajoie, Z.P. Fan, A.A. Sigova, J. Reddy, D. Borges-Rivera, T.I. Lee, R. Jaenisch, M.H. Porteus, J. Dekker, and R.A. Young, Activation of proto-oncogenes by disruption of chromosome neighborhoods. Science, 2016. 351(6280): p. 1454–1458.

16. Narendra, V., M. Bulajic, J. Dekker, E.O. Mazzoni, and D. Reinberg, CTCF-mediated topological boundaries during development foster appropriate gene regulation. Genes Dev, 2016. 30(24): p. 2657–2662.

17. Merkenschlager, M. and E.P. Nora, CTCF and Cohesin in genome folding and transcriptional gene regulation. Annu Rev Genomics Hum Genet, 2016. 17: p. 17–43.

18. Sanborn, A.L., S.S.P. Rao, S.C. Huang, N.C. Durand, M.H. Huntley, A.I. Jewett, I.D. Bochkov, D. Chinnappan, A. Cutkosky, J. Li, K.P. Geeting, A. Gnirke, A. Melnikov, D. McKenna, E.K. Stamenova, E.S. Lander, and E.L. Aiden, Chromatin extrusion explains key features of loop and domain formation in wild-type and engineered genomes. Proc Natl Acad Sci U S A, 2015. 112(47): p. E6456–E6465.

19. Fudenberg, G., M. Imakaev, C. Lu, A. Goloborodko, N. Abdennur, and L.A. Mirny, Formation of chromosomal domains by loop extrusion. Cell Rep, 2016. 15(9): p. 2038–2049.

20. Nuebler, J., G. Fudenberg, M. Imakaev, N. Abdennur, and L.A. Mirny, Chromatin organization by an interplay of loop extrusion and compartmental segregation. Proc Natl Acad Sci U S A, 2018. 115(29): p. E6697–E6706.

21. Tang, Z., O.J. Luo, X. Li, M. Zheng, J.J. Zhu, P. Szalaj, P. Trzaskoma, A. Magalska, J. Wlodarczyk, B. Ruszczycki, P. Michalski, E. Piecuch, P. Wang, D. Wang, S.Z. Tian, M. Penrad-Mobayed, L.M. Sachs, X. Ruan, C.L. Wei, E.T. Liu, G.M. Wilczynski, D. Plewczynski, G. Li, and Y. Ruan, CTCF-mediated human 3D genome architecture reveals chromatin topology for transcription. Cell, 2015. 163(7): p. 1611–1627.

22. Lupianez, D.G., K. Kraft, V. Heinrich, P. Krawitz, F. Brancati, E. Klopocki, D. Hom, H. Kayserili, J.M. Opitz, R. Laxova, F. Santos-Simarro, B. Gilbert-Dussardier, L. Wittler, M. Borschiwer, S.A. Haas, M. Osterwalder, M. Franke, B. Timmermann, J. Hecht, M. Spielmann, A. Visel, and S. Mundlos, Disruptions of Topological Chromatin Domains Cause Pathogenic Rewiring of Gene-Enhancer Interactions. Cell, 2015. 161(5): p. 1012–1025.

23. Hou, C., H. Zhao, K. Tanimoto, and A. Dean, CTCF-dependent enhancer-blocking by alternative chromatin loop formation. Proc Natl Acad Sci U S A, 2008. 105(51): p. 20398–20403.

24. Flavahan, W.A., Y. Drier, B.B. Liau, S.M. Gillespie, A.S. Venteicher, A.O. Stemmer-Rachamimov, M.L. Suva, and B.E. Bernstein, Insulator dysfunction and oncogene activation in IDH mutant gliomas. Nature, 2016. 529(7584): p. 110–114.

25. Wu, Q. and T. Maniatis, A striking organization of a large family of human neural cadherin-like cell adhesion genes. Cell, 1999. 97(6): p. 779–790.

26. Lefebvre, J.L., D. Kostadinov, W.V. Chen, T. Maniatis, and J.R. Sanes, Protocadherins mediate dendritic self-avoidance in the mammalian nervous system. Nature, 2012. 488(7412): p. 517–521.

27. Toyoda, S., M. Kawaguchi, T. Kobayashi, E. Tarusawa, T. Toyama, M. Okano, M. Oda, H. Nakauchi, Y. Yoshimura, M. Sanbo, M. Hirabayashi, T. Hirayama, T. Hirabayashi, and T. Yagi, Developmental epigenetic modification regulates stochastic expression of clustered protocadherin genes, generating single neuron diversity. Neuron, 2014. 82(1): p. 94–108.

28. Schreiner, D. and J.A. Weiner, Combinatorial homophilic interaction between gamma-protocadherin multimers greatly expands the molecular diversity of cell adhesion. Proc Natl Acad Sci U S A, 2010. 107(33): p. 14893–14898.

29. Chen, W.V., C.L. Nwakeze, C.A. Denny, S. O’Keeffe, M.A. Rieger, G. Mountoufaris, A. Kirner, J.D. Dougherty, R. Hen, Q. Wu, and T. Maniatis, Pcdhalphac2 is required for axonal tiling and assembly of serotonergic circuitries in mice. Science, 2017. 356(6336): p. 406–411.

30. Fan, L., Y.C. Lu, X.L. Shen, H. Shao, L. Suo, and Q. Wu, Alpha protocadherins and Pyk2 kinase regulate cortical neuron migration and cytoskeletal dynamics via Rac1 GTPase and WAVE complex in mice. Elife, 2018. 7.

31. Mountoufaris, G., D. Canzio, C.L. Nwakeze, W.V. Chen, and T. Maniatis, Writing, Reading, and Translating the Clustered Protocadherin Cell Surface Recognition Code for Neural Circuit Assembly. Annu Rev Cell Dev Biol, 2018. 34: p. 471–493.

32. Jain, S., Z. Ba, Y. Zhang, H.Q. Dai, and F.W. Alt, CTCF-binding elements mediate accessibility of RAG substrates during chromatin scanning. Cell, 2018.

33. Kehayova, P., K. Monahan, W. Chen, and T. Maniatis, Regulatory elements required for the activation and repression of the protocadherin-alpha gene cluster. Proc Natl Acad Sci U S A, 2011. 108(41): p. 17195–17200.

34. Guo, Y., K. Monahan, H. Wu, J. Gertz, K.E. Varley, W. Li, R.M. Myers, T. Maniatis, and Q. Wu, CTCF/cohesin-mediated DNA looping is required for protocadherin alpha promoter choice. Proc Natl Acad Sci U S A, 2012. 109(51): p. 21081–21086.

35. Allahyar, A., C. Vermeulen, B.A.M. Bouwman, P.H.L. Krijger, M. Verstegen, G. Geeven, M. van Kranenburg, M. Pieterse, R. Straver, J.H.I. Haarhuis, K. Jalink, H. Teunissen, I.J. Renkens, W.P. Kloosterman, B.D. Rowland, E. de Wit, J. de Ridder, and W. de Laat, Enhancer hubs and loop collisions identified from single-allele topologies. Nat Genet, 2018. 50(8): p. 1151–1160.

36. Esumi, S., N. Kakazu, Y. Taguchi, T. Hirayama, A. Sasaki, T. Hirabayashi, T. Koide, T. Kitsukawa, S. Hamada, and T. Yagi, Monoallelic yet combinatorial expression of variable exons of the protocadherin-alpha gene cluster in single neurons. Nat Genet, 2005. 37(2): p. 171–176.

37. Tasic, B., Z. Yao, L.T. Graybuck, K.A. Smith, T.N. Nguyen, D. Bertagnolli, J. Goldy, E. Garren, M.N. Economo, S. Viswanathan, O. Penn, T. Bakken, V. Menon, J. Miller, O. Fong, K.E. Hirokawa, K. Lathia, C. Rimorin, M. Tieu, R. Larsen, T. Casper, E. Barkan, M. Kroll, S. Parry, N.V. Shapovalova, D. Hirschstein, J. Pendergraft, H.A. Sullivan, T.K. Kim, A. Szafer, N. Dee, P. Groblewski, I. Wickersham, A. Cetin, J.A. Harris, B.P. Levi, S.M. Sunkin, L. Madisen, T.L. Daigle, L. Looger, A. Bernard, J. Phillips, E. Lein, M. Hawrylycz, K. Svoboda, A.R. Jones, C. Koch, and H. Zeng, Shared and distinct transcriptomic cell types across neocortical areas. Nature, 2018. 563(7729): p. 72–78.

38. Shou, J., J. Li, Y. Liu, and Q. Wu, Precise and predictable CRISPR chromosomal rearrangements reveal principles of Cas9-mediated nucleotide insertion. Mol Cell, 2018. 71: p. 498–509.

39. Srinivasan, M., J.C. Scheinost, N.J. Petela, T.G. Gligoris, M. Wissler, S. Ogushi, J.E. Collier, M. Voulgaris, A. Kurze, K.L. Chan, B. Hu, V. Costanzo, and K.A. Nasmyth, The Cohesin Ring Uses Its Hinge to Organize DNA Using Non-topological as well as Topological Mechanisms. Cell, 2018. 173(6): p. 1508–1519 e1518.

40. ENCODE Project Consortium, An integrated encyclopedia of DNA elements in the human genome. Nature, 2012. 489(7414): p. 57–74.

41. Canzio, D., C. Nwakeze, A. Horta, S. Rajkumar, E. Coffey, E. Duffy, R. Duffié, M. Simon, S. Lomvardas, and T. Maniatis, Antisense lncRNA transcription drives stochastic protocadherin α promoter choice. bioRxiv, 2018.

42. Wu, Q., T. Zhang, J.F. Cheng, Y. Kim, J. Grimwood, J. Schmutz, M. Dickson, J.P. Noonan, M.Q. Zhang, R.M. Myers, and T. Maniatis, Comparative DNA sequence analysis of mouse and human protocadherin gene clusters. Genome Res, 2001. 11(3): p. 389–404.

43. Tanimoto, K., Q. Liu, J. Bungert, and J.D. Engel, Effects of altered gene order or orientation of the locus control region on human beta-globin gene expression in mice. Nature, 1999. 398(6725): p. 344–348.

44. Busslinger, G.A., R.R. Stocsits, P. van der Lelij, E. Axelsson, A. Tedeschi, N. Galjart, and J.M. Peters, Cohesin is positioned in mammalian genomes by transcription, CTCF and Wapl. Nature, 2017. 544(7651): p. 503–507.

45. Li, J., J. Shou, Y. Guo, Y. Tang, Y. Wu, Z. Jia, Y. Zhai, Z. Chen, Q. Xu, and Q. Wu, Efficient inversions and duplications of mammalian regulatory DNA elements and gene clusters by CRISPR/Cas9. J Mol Cell Biol, 2015. 7(4): p. 284–298.

46. Vian, L., A. Pekowska, S.S.P. Rao, K.R. Kieffer-Kwon, S. Jung, L. Baranello, S.C. Huang, L. El Khattabi, M. Dose, N. Pruett, A.L. Sanborn, A. Canela, Y. Maman, A. Oksanen, W. Resch, X.W. Li, B. Lee, A.L. Kovalchuk, Z.H. Tang, S. Nelson, M. Di Pierro, R.R. Cheng, I. Machol, B.G. St Hilaire, N.C. Durand, M.S. Shamim, E.K. Stamenova, J.N. Onuchic, Y.J. Ruan, A. Nussenzweig, D. Levens, E.L. Aiden, and R. Casellas, The Energetics and Physiological Impact of Cohesin Extrusion. Cell, 2018. 173(5): p. 1165–+.

47. Dostie, J. and J. Dekker, Mapping networks of physical interactions between genomic elements using 5C technology. Nat Protoc, 2007. 2(4): p. 988–1002.

48. Dempster, A.P., N.M. Laird, and D.B. Rubin, Maximum likelihood from incomplete data via the EM algorithm. Journal of the Royal Statistical Society. Series B (Methodological), 1977. 39(1): p. 1–38.

49. Zhang, N., S.G. Kuznetsov, S.K. Sharan, K. Li, P.H. Rao, and D. Pati, A handcuff model for the cohesin complex. J Cell Biol, 2008. 183(6): p. 1019–1031.

50. Ganji, M., I.A. Shaltiel, S. Bisht, E. Kim, A. Kalichava, C.H. Haering, and C. Dekker, Real-time imaging of DNA loop extrusion by condensin. Science, 2018. 360(6384): p. 102–105.

51. Alipour, E. and J.F. Marko, Self-organization of domain structures by DNA-loop-extruding enzymes. Nucleic Acids Res, 2012. 40(22): p. 11202–11212.

52. Yin, M.L., J.Y. Wang, M. Wang, X.M. Li, M. Zhang, Q. Wu, and Y.L. Wang, Molecular mechanism of directional CTCF recognition of a diverse range of genomic sites. Cell Research, 2017. 27(11): p. 1365–1377.

